# An interpretable bimodal neural network characterizes the sequence and preexisting chromatin predictors of induced TF binding

**DOI:** 10.1101/672790

**Authors:** Divyanshi Srivastava, Begüm Aydin, Esteban O. Mazzoni, Shaun Mahony

**Affiliations:** Center for Eukaryotic Gene Regulation, Department of Biochemistry & Molecular Biology, Pennsylvania State University, University Park, PA; Department of Biology, New York University, New York, NY

## Abstract

Transcription factor (TF) binding specificity is determined via a complex interplay between the TF’s DNA binding preference and cell type-specific chromatin environments. The chromatin features that correlate with TF binding in a given cell type have been well characterized. For instance, the binding sites for a majority of TFs display concurrent chromatin accessibility. However, concurrent chromatin features reflect the binding activities of the TF itself, and thus provide limited insight into how genome-wide TF-DNA binding patterns became established in the first place. To understand the determinants of TF binding specificity, we therefore need to examine how newly activated TFs interact with sequence and preexisting chromatin landscapes.

Here, we investigate the sequence and preexisting chromatin predictors of TF-DNA binding by examining the genome-wide occupancy of TFs that have been induced in well-characterized chromatin environments. We develop Bichrom, a bimodal neural network that jointly models sequence and preexisting chromatin data to interpret the genome-wide binding patterns of induced TFs. We find that the preexisting chromatin landscape is a differential global predictor of TF-DNA binding; incorporating preexisting chromatin features improves our ability to explain the binding specificity of some TFs substantially, but not others. Furthermore, by analyzing site-level predictors, we show that TF binding in previously inaccessible chromatin tends to correspond to the presence of more favorable cognate DNA sequences. Bichrom thus provides a framework for modeling, interpreting, and visualizing the joint sequence and chromatin landscapes that determine TF-DNA binding dynamics.

## Introduction

Sequence-specific transcription factors (TFs) bind DNA at their cognate sequence motifs, using both direct base interactions and DNA structural feature recognition^1–3^. However, the presence of cognate motif instances alone is a poor predictor of TF binding^4,5^. TFs typically bind a small fraction of their potential target motif instances in a given cell type, and the cohort of sites which are bound can vary greatly across cell types^6–8^. These observations suggest that cell type-specific TF selectivity is governed by cell type-specific chromatin environments^5,7,9,10^. Cell-specific patterns of chromatin accessibility, nucleosome positioning, and histone post-translational modifications modify the availability of a TF’s sequence motifs^11–16^. Co-operative interactions with other regulatory proteins can alter a TF’s intrinsic sequence preferences, or enable binding at otherwise unavailable target sequence motifs^5,17–19^. Even pioneer TFs, which are characterized by their ability to bind target motifs in relatively inaccessible chromatin, bind DNA in cell type-specific patterns that can be modulated by other TFs^20–23^. Thus, it remains unclear how DNA sequence, chromatin structure, and interactions with other regulators act in concert to determine cell type-specific TF binding patterns.

Most previous searches for features associated with cell type-specific TF binding sites have performed correlations with chromatin data measured when the TFs under study are already bound to DNA (i.e., “concurrent” chromatin information)^24–28^. But TFs and their recruited regulatory complexes often alter local chromatin landscapes^29,30^. Concurrent chromatin features thus cannot be used to address the question of how TF binding patterns become established in the first place. The few studies that have analyzed the determinants of TF binding in dynamic contexts have lacked integrated analysis approaches that separate DNA sequence and prior chromatin predictors of future TF binding activities^9,11,12,21,31,32^.

We present Bichrom, a bimodal neural network framework for characterizing the relative contributions of DNA sequence and preexisting cell type-specific chromatin landscape to an induced TF’s binding specificity (**Fig. 1A**). Our use of neural networks is motivated by their advantages in predicting genome-wide TF binding patterns^24,28,33^, and the ability of multi-modal neural networks to integrate heterogeneous data types^34–36^. Bichrom’s architecture embeds TF binding sites into a two-dimensional latent space, which can be used to estimate the relative contributions of DNA sequence and preexisting chromatin features at individual TF binding sites. By comparing how well neural networks can represent genome-wide binding patterns using sequence information alone versus a combination of sequence and preexisting chromatin features, we can quantify the marginal amount of information added by preexisting chromatin. Comparing such metrics across TFs allows us to assess how TFs differ in their overall sensitivity to preexisting chromatin.

**Figure 1:**
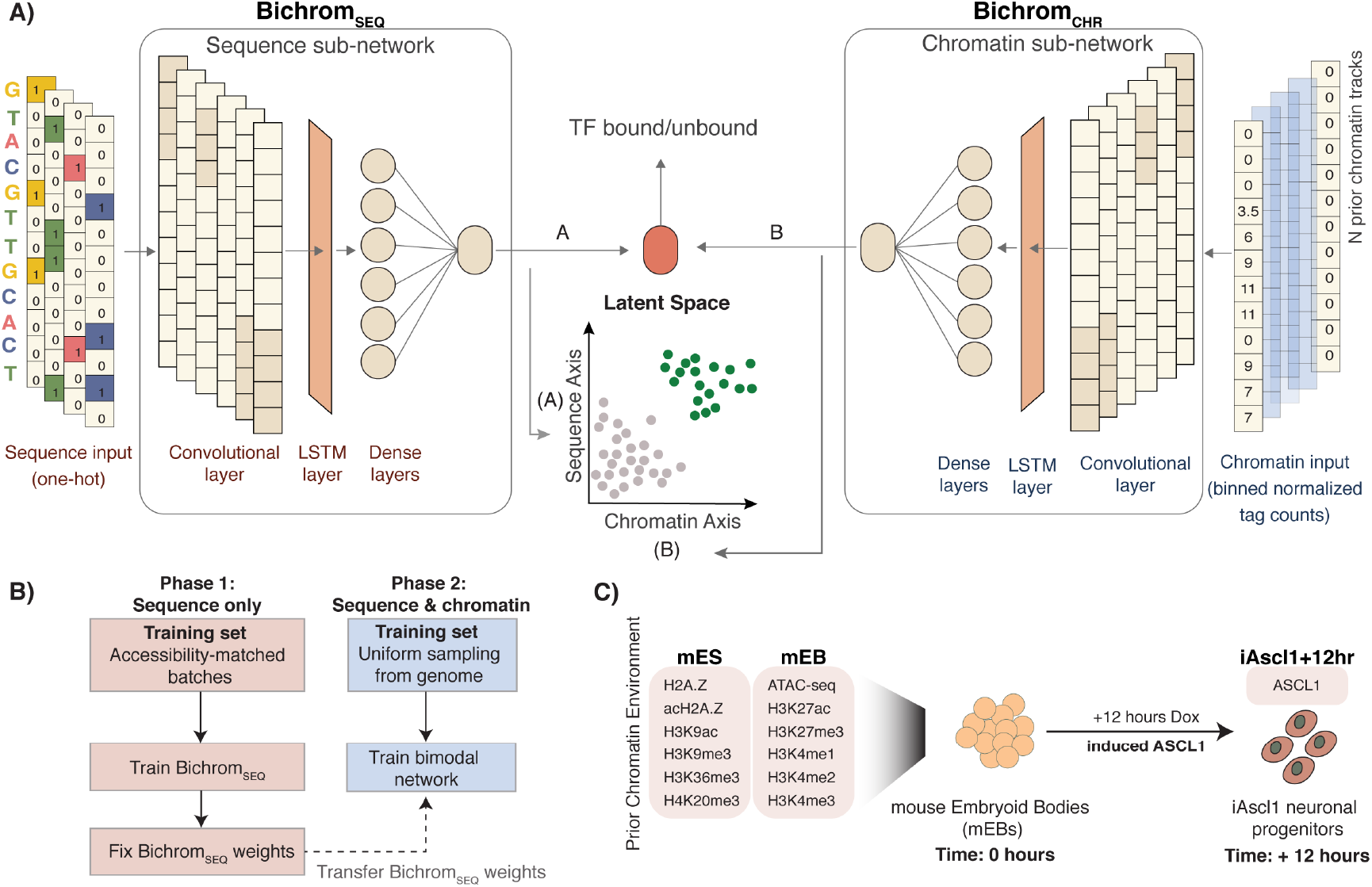
**A)** Bichrom’s bimodal sequence and preexisting chromatin network consists of two sub-networks: the sequence sub-network (Bichrom_SEQ_) which uses one-hot encoded DNA sequence as input; and the chromatin subnetwork (Bichrom_CHR_) which uses binned normalized tag counts from chromatin experiments such as ATAC-seq and histone modification ChIP-seq. The sequence and chromatin sub-network activations embed the training data into a lower-dimensional plane, which is then used by a sigmoid-activated node for TF binding label classification (i.e., bound/unbound). **B)** Overview of Bichrom’s training strategy. Bichrom_SEQ_ is trained using training batches within which positive and negative training samples are matched in their prior accessibility status. The weights of the convolutional and LSTM layers of Bichrom_SEQ_ are fixed, and Bichrom is trained using both sequence and preexisting chromatin data. **C)** Overview of the Ascl1 data: Ascl1 expression is induced in mouse embryoid bodies (mEBs) using a Dox-inducible promoter and Ascl1 binding is measured 12 hours post induction. Bichrom training uses 12 prior chromatin datasets from mEB and mES cell types.

Our approach is distinct from other recent applications of neural networks to TF binding prediction tasks. Several studies have also used neural networks to integrate DNA sequence and concurrent chromatin landscape information, but with the goal of imputing unobserved TF binding patterns in a given cell type^24,28,37^. In contrast, we focus on settings in which we already know (via ChIP-seq) where a TF of interest is binding when expressed in a given cell type, and we aim to interpret how those binding sites relate to DNA sequence and preexisting chromatin features. Previous applications that impute TF binding have also used early-integration frameworks (i.e., DNA sequence and chromatin accessibility data are integrated at a feature level), thereby making it challenging to interpret the contributions of individual features towards TF binding at individual sites^6,24,26,37–39^. Our bimodal architecture has the distinct advantage of enabling deconvolution of sequence and chromatin predictors of TF-DNA binding.

We demonstrate Bichrom’s utility by examining the binding determinants of the pro-neural bHLH TF Ascl1 when it is induced in mouse embryoid body (mEB) cells^40^. Bichrom analysis finds that despite Ascl1’s characterized pioneering abilities^40–44^, Ascl1 binding is dependent on the preexisting mEB chromatin environment at a subset of its binding sites. Next, we demonstrate that Bichrom can informatively rank TFs by their relative dependence on prior chromatin landscapes. We focus on a selection of neuronal TFs that are activated downstream of Ascl1 (and bind DNA in the chromatin environment established by Ascl1)^40^; Bichrom’s assessment of relative chromatin dependence for these TFs is supported by observing whether the same TFs bind different sites when they become activated in a different chromatin environment. Finally, we expand our analysis to examine the differential sequence and prior chromatin drivers for 12 TFs induced in cell types for which the preexisting chromatin accessibility landscape has also been characterized (mouse and human fibroblasts^23,45^). While we focus here on systems in which TF expression is induced in cell lines, Bichrom is broadly applicable to study TF binding determinants in any dynamic regulatory system in which chromatin landscapes can be assayed before a TF’s DNA binding activity occurs.

## Results

### A bimodal neural network integrates DNA sequence and preexisting chromatin data to characterize TF binding predictors

Bichrom’s bimodal neural network is composed of two sub-networks: one that operates only on DNA sequence features (Bichrom_SEQ_), and one that operates on chromatin features derived from ATAC-seq and ChIP-seq specific for histone modifications (Bichrom_CHR_). Both sub-networks are designed to output real-valued activations, which are additively combined by a single sigmoid-activated node (**Fig. 1A**). Bichrom is trained to predict TF binding labels, as defined by ChIP-seq peaks. In our typical usage, Bichrom trains on binding labels for a TF that has become activated in a given cell type, using input features from DNA sequence and chromatin data profiled before the targeted TF has become active. While we phrase Bichrom training as a predictive task, our motivation is not to predict TF binding (which, of course, has already been observed via ChIP-seq), but rather to characterize the sequence and preexisting chromatin states that define a TF’s binding pattern. We do so by taking advantage of the interpretable nature of the bimodal network architecture; the weighted activations flowing from each sub-network result in a latent two-dimensional representation of each genomic region, which we can analyze to interpret and compare the sequence and preexisting chromatin predictors at individual TF binding sites (**Fig. 1A**).

Bichrom’s neural network hyper-parameters were chosen via a random grid-search (see **Methods**). Each sub-network consists of a single convolutional neural network (CNN) layer that acts as a primary feature extractor, followed by a long short-term memory (LSTM) layer that can capture potential interactions between convolutional filters^24,46^. The LSTM output is fed through dense layers, which are combined to produce a scalar real-valued output (**Fig. 1A**). Bichrom’s CNN-LSTM architecture was found to perform better or equivalent to a wide range of alternative CNN-based architectures (see **Methods**, **Supp. Fig. 1**). Each sub-network is input features from 500bp genomic windows: Bichrom_SEQ_ operates on one-hot encoded DNA sequences, while Bichrom_CHR_ operates on normalized and binned read counts from one or more preexisting chromatin data tracks. Since we aim to interpret the sub-network activations as separable sources of sequence and prior chromatin information, we wish to avoid scenarios where the Bichrom_SEQ_ sub-network learns sequence features that are associated with the prior chromatin landscape. To minimize such confounding issues, we train Bichrom_SEQ_ using mini-batches where the positive and negative sets are matched in their prior accessibility labels (see **Methods, Fig. 1B**).

While our primary motivation is the interpretation of sequence and prior chromatin predictors of induced TF binding, we asked whether Bichrom’s architecture and training scheme produces an accurate representation of genome-wide TF-DNA binding data. To ensure that Bichrom can represent genome-wide TF binding patterns with accuracies approaching current state-of-the-art predictive methods, we assessed Bichrom’s performance on within-cell-type predictive tasks from the ENCODE-DREAM challenge^24,26,28,47^. This challenge evaluated the ability of various methods to predict genome-wide binding of several human TFs given concurrent chromatin accessibility (DNase-seq) and gene expression data. Using only sequence and concurrent chromatin accessibility data, Bichrom predicts TF-DNA binding in held-out genomic regions with accuracies comparable to, albeit slightly lower than, the top models described in the challenge (see **Methods, Supp. Fig. 2**). Thus, Bichrom’s bimodal architecture does not negatively impact its ability to represent genome-wide TF-DNA binding patterns using sequence and chromatin features.

### Ascl1 binding sites are partially predicted by preexisting chromatin information

Ascl1 is a proposed pioneer TF which can bind to relatively inaccessible sites when induced in fibroblasts and early embryonic cell types^40–44^. Whether the preexisting chromatin environment plays a role in determining induced Ascl1 binding sites remains unclear^42^. To address this question, we applied Bichrom to interpret the relationships between induced Ascl1 binding and preexisting sequence and chromatin landscapes. Specifically, we train Bichrom using previously published Ascl1 ChIP-seq data measured 12 hours after Ascl1 expression has been induced in mouse embryoid body (mEB) cells^40^ (**Fig. 1C**). The 12 hour timepoint is the earliest at which we can obtain robust Ascl1 ChIP-seq binding data, and is thus the most likely to represent the initial binding activities that are shaped by previous chromatin states. In this analysis, the Bichrom_CHR_ sub-network is trained using 12 chromatin-related datasets from mEB and related mouse embryonic stem (mES) cells: ATAC-seq, H2A.Z, acH2A.Z, H3K27ac, H3K27me3, H3K4me1/me2/me3, H3K9ac, H3K9me3, H3K36me3, and H4K20me3 (**Supp. Tables 1 and 2, Fig. 1C**).

To assess whether the preexisting chromatin landscape is predictive of future Ascl1 binding locations, we compared Bichrom performance with a baseline neural network trained only with sequence information. The baseline sequence-only network was constructed using the same hyper-parameters and architecture as the Bichrom_SEQ_ sub-network. Training was repeated 9 times for each network, each training round using a separate held-out test chromosome. The sequence-only network predicts induced Ascl1 binding with a median area under the precision-recall curve (auPRC) of 0.42. In contrast, Bichrom predicts induced Ascl1 binding with a median auPRC of 0.59, suggesting that information in the preexisting chromatin landscape significantly improves prediction of Ascl1 binding (Wilcoxon signed rank test p-value 0.003, **Fig. 2A**). As a negative control, Bichrom trained using sequence and a ChIP input control experiment instead of preexisting chromatin data does not lead to significant improvement in network performance when compared to a sequence-only network (auPRC=0.45, **Fig. 2A**). Additionally, we confirmed that Bichrom’s additive bimodal design does not perform worse than a model with more complex interactions between sequence and prior chromatin features (**Suppl. Fig. 3B**).

**Figure 2:**
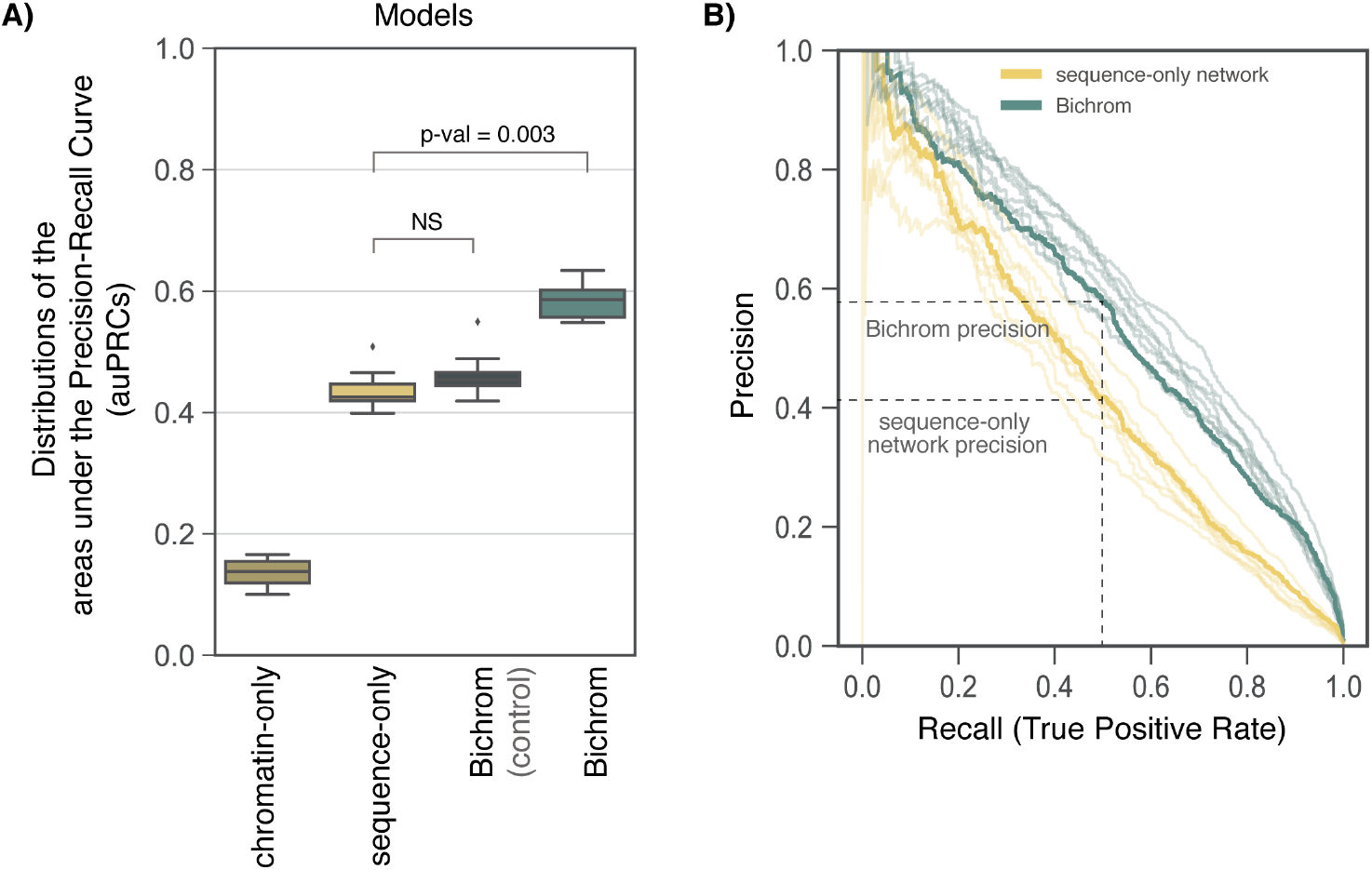
**A)** Distribution of model performance (auPRC) in predicting Ascl1 binding sites for a neural network trained using only chromatin data (chromatin-only), a neural network trained using only sequence data (sequence-only), Bichrom using sequence and a ChIP input control experiment (Bichrom control), and Bichrom using sequence and 12 preexisting ES chromatin datasets. The boxplots represent data from 9 independent training sets, each consisting of a separate held-out test chromosome. **B)** The precision-recall curves for 9 models, each tested on 9 distinct held-out test chromosomes. The P-R curves for networks that use a training set in which chromosome 10 is held-out for testing are highlighted in solid lines, performance for other training sets is represented with lighter (alpha=0.2) traces. Precision at a fixed recall of 0.5 is highlighted for both the sequence-only network and Bichrom.

Notably, the improved performance of Bichrom’s bimodal network is driven largely by improved specificity. At a false positive rate of 0.05, a majority of Ascl1-bound sites are correctly predicted by both the sequence-only network and Bichrom (**Suppl. Fig. 3A**). However, at a fixed recall of 0.5, Bichrom’s precision is substantially greater than that of the sequence-only network (**Fig. 2B**). Thus, the incorporation of mEB and mES chromatin information significantly improves Bichrom’s ability to predict induced Ascl1 binding in a future timepoint, suggesting that although it is an established pioneer TF, Ascl1 binding sites are partially determined by the preexisting chromatin landscape.

### Bichrom deconvolves the sequence and preexisting chromatin predictors of induced Ascl1 binding

Beyond quantifying preexisting chromatin’s overall contribution to improving Ascl1 binding predictions, Bichrom’s unique bimodal architecture enables decomposition of sequence and prior chromatin predictors at individual Ascl1 binding sites. For a given genomic window, Bichrom’s TF binding score is a simple linear combination of the activations from Bichrom_SEQ_ and Bichrom_CHR_ sub-networks. Thus, every genomic window can be embedded in a two-dimensional latent space defined by the sub-network activations, enabling an intuitive visualization of how much the sequence and prior chromatin sub-networks contributed to the overall predictive score (**Fig. 1A**).

Applied to the Bichrom network trained on Ascl1 binding data, we find that Ascl1-bound genomic windows (orange) are well-separated from randomly sampled unbound genomic windows (grey) in the twodimensional latent space (**Fig. 3A**). However, Ascl1 binding sites are distributed over a broad range of Bichrom_SEQ_ and Bichrom_CHR_ sub-network activation scores. The variation in sub-network activations suggests that some of Bichrom’s Ascl1 binding site predictions are driven primarily by sequence information, while some are driven by preexisting chromatin features (**Fig. 3A**). Furthermore, the diversity in Ascl1 sequence sub-network (Bichrom_SEQ_) scores is higher at sites that are scored favorably by the Bichrom_CHR_ chromatin sub-network, suggesting that the presence of a favorable preexisting chromatin environment may enable Ascl1 binding even at sites with relatively weaker sequence features (**Fig. 3B**). We confirm that this compensatory effect is not an artifact of the linear combination between the sequence and chromatin subnetworks by fixing the sequence sub-network weights while training the chromatin sub-network (see **Methods**, **Supp. Fig. 5**). Thus, Bichrom learns a model in which Ascl1 binding sites exhibit a broad range of sequence and chromatin sub-network scores, where favorable prior chromatin features can partially compensate for weaker sequence features.

**Figure 3:**
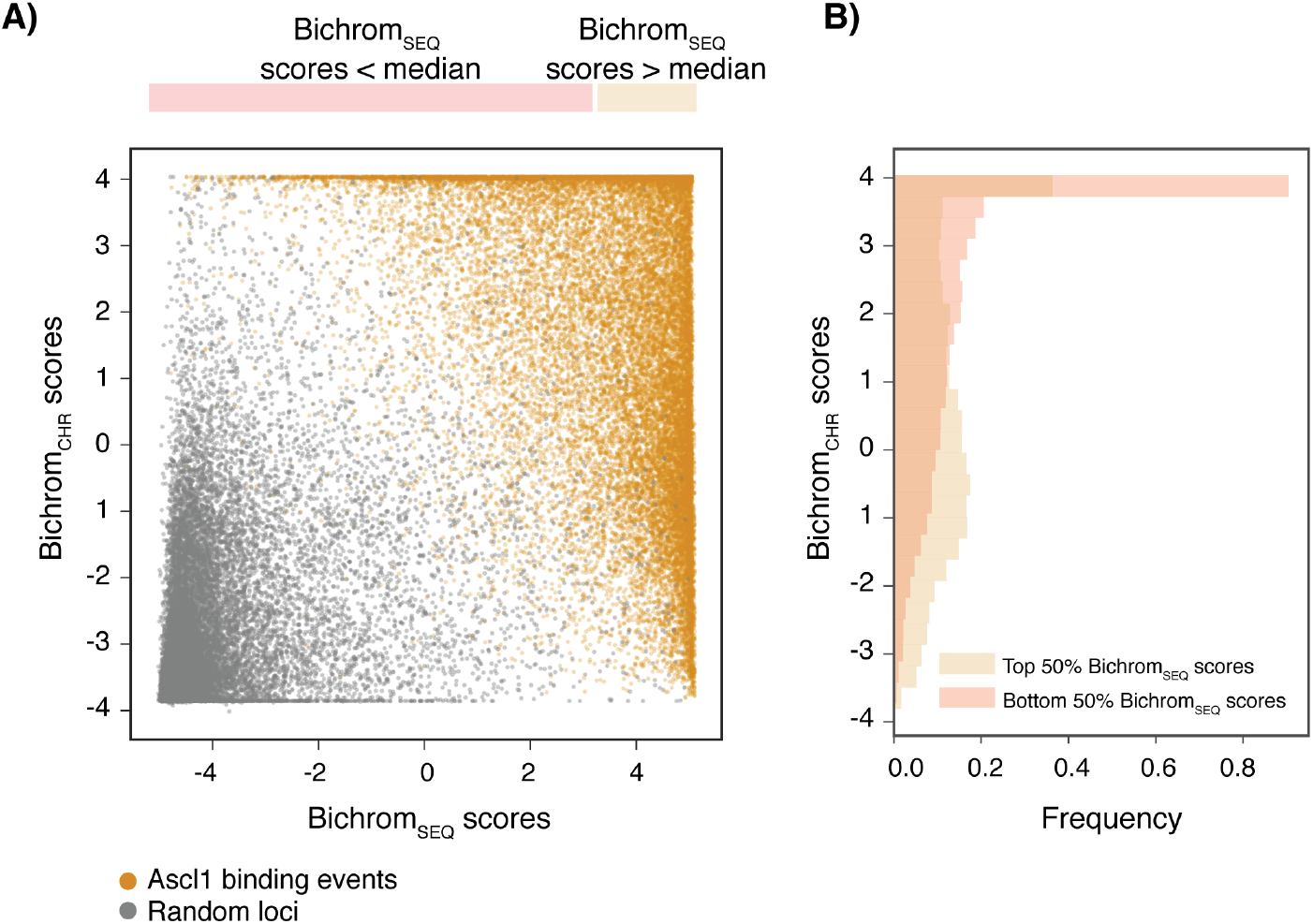
**A)** Bichrom network-derived latent representation of induced Ascl1 binding events (orange) and a randomly sampled subset of unbound genomic regions (grey). The axes represent contributions of the Bichrom_SEQ_ sub-network and the Bichrom_CHR_ sub-network toward bimodal network predictions. **B)** Distributions of Bichrom_CHR_ scores for Ascl1 binding events conditioned on the Bichrom_SEQ_ scores.

To investigate informative sequence features at Ascl1 binding sites, we used an integrated gradients^48^ feature attribution-based approach to identify local sequence windows driving high Bichrom_SEQ_ sub-network scores (see **Methods, Supp. Fig. 6A, B**). As expected, regions with high Bichrom_SEQ_ scores contain motifs related to the CAGSTG E-box, consistent with Ascl1’s cognate DNA binding preference^40^ (**Fig. 4A**). We also find a POU homeodomain DNA binding motif, suggesting that Ascl1 may bind a subset of its sites either in concert with, or at sites pre-bound by, a POU domain TF such as Oct4 (one of the main regulators of pluripotency in the preexisting embryonic cells) (**Fig. 4A**). Bichrom_SEQ_ scores are strongly associated with Ascl1 cognate motif multiplicity; Ascl1 binding sites that receive high Bichrom_SEQ_ scores contain higher frequencies of E-box motif instances (**Supp. Fig. 6C**) and CAGSTG *k*-mers (**Fig. 4B, C**). The correlation between motif multiplicity and Bichrom_SEQ_ scores is maintained when scoring randomly sampled sequences in which we inserted variable numbers of CAGSTG *k*-mer instances (**Fig. 4D**). We also investigated whether Bichrom_SEQ_ scores encapsulate subtle nucleotide composition dependences in regions flanking Ascl1’s cognate binding motif^2,40,49^. Specifically, we inserted CAGSTG *k*-mers flanked by variable nucleotides into an artificial uniform sequence (see **Methods**), and found that Bichrom_SEQ_ scores vary substantially according to specific motif-flanking nucleotide compositions (**Fig. 4E**).

**Figure 4:**
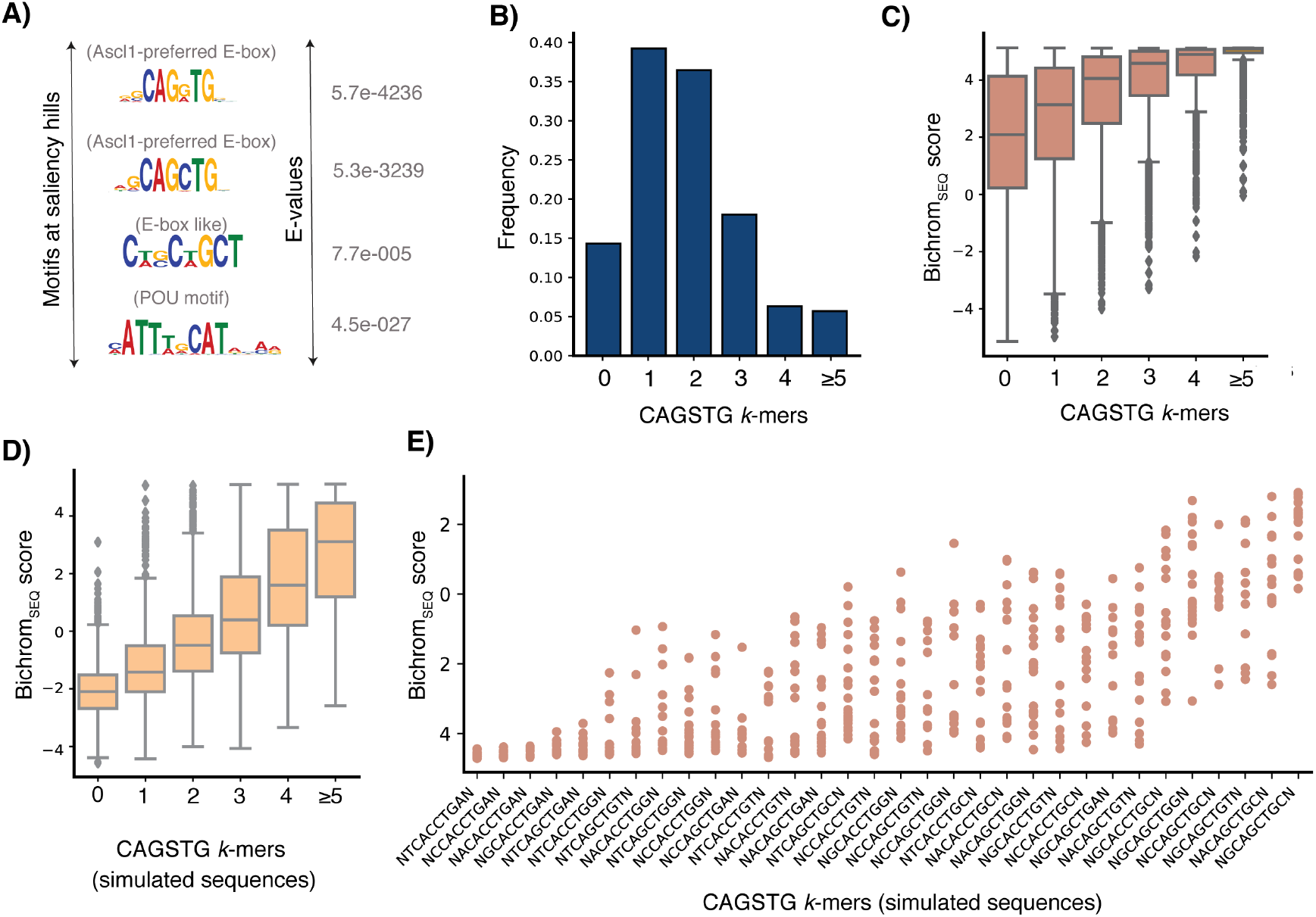
**A)** Motifs enriched at saliency hills (regions of the input sequences that are used by the sequence sub-network to make predictions). **B)** Frequencies of CAGSTG *k*-mers at all Ascl1 binding sites. **C)** Bichrom_SEQ_ scores increase with increasing motif multiplicity at Ascl1 binding windows. **D)** Embedding CAGSTG motifs in simulated sequences confirms that the Bichrom_SEQ_ sub-network uses the number of motif occurrences as a score-driving predictor. **E)** Bichrom_SEQ_ scores assigned to CAGSTG *k*-mers vary widely based on nucleotides flanking the cognate *k*-mer.

We next investigated the preexisting chromatin features driving variability in Bichrom_CHR_ sub-network scores. Ascl1 binding sites that receive the highest Bichrom_CHR_ scores are enriched in chromatin signals associated with regulatory activity in the preexisting mES and mEB cell types (e.g. preexisting ATAC-seq, H3K27ac, H3K4me2, H3K4me3, H3K9ac; **Fig. 5A**). Further, genome-wide domains of preexisting ATAC-seq and “active” histone mark ChIP-seq enrichment are more likely to receive higher Bichrom_CHR_ scores (**Fig. 5B**). H3K9me3 and H4K20me3 domains receive the lowest Bichrom_CHR_ scores, in contrast to a previous suggestion that preexisting H3K9me3 is predictive of induced Ascl1 binding in mouse embryonic fibroblasts^42^ (**Fig. 5B**).

**Figure 5:**
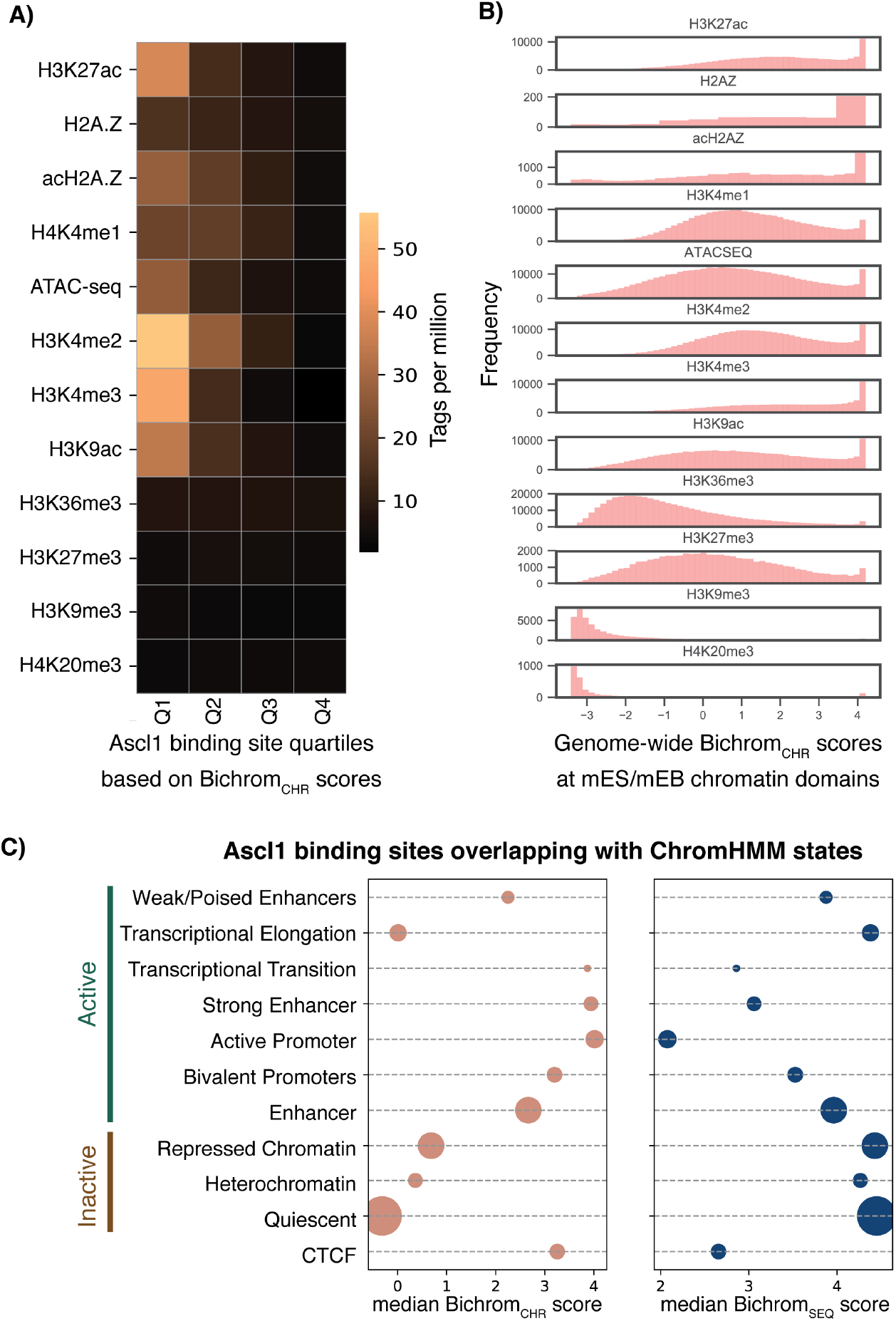
**A)** Mean mES/mEB chromatin feature tag enrichment at Ascl1 binding sites, divided into quartiles based on their association with Bichrom_CHR_ sub-network scores. Sites belonging to the highest-scoring quartile are enriched for preexisting active histone modifications such as H3K4me2, H3K4me3, H3K27ac, and H3K9ac. The lowest-scoring quartile lack enrichment of any measured histone modifications. **B)** The distribution of Bichrom_CHR_ sub-network scores at genomic regions enriched for each of the mES/mEB chromatin features. **C)** Median Bichrom_SEQ_ and Bichrom_CHR_ sub-network scores at Ascl1 sites that overlap mES/mEB ChromHMM states. Bubble size corresponds to relative proportion of Ascl1 binding sites. Median Bichrom_SEQ_ sub-network scores are typically lower at states with high Bichrom_CHR_ sub-network scores.

Consistent with our analyses of individual chromatin signals, Ascl1 binding sites that overlap various ChromHMM-defined^50^ mES/mEB chromatin states receive widely varying median Bichrom_CHR_ scores (**Fig. 5C**). Ascl1 binding sites that overlap preexisting active promoter and strong enhancer states receive the highest Bichrom_CHR_ scores, while Polycomb-repressed (marked by H3K27me3), heterochromatin (marked by H3K9me3), and quiescent states receive the lowest Bichrom_CHR_ scores (**Fig. 5C**). In accordance with our observations from the latent space embeddings (**Fig. 3**), Ascl1 binding sites located in preexisting enhancer or promoter states receive lower median scores from the Bichrom_SEQ_ sequence sub-network (**Fig. 5C**). Conversely, Ascl1 sites in preexisting heterochromatin and quiescent states receive the highest median Bichrom_SEQ_ scores.

Taken together, our Bichrom analysis of induced Ascl1 binding sites suggest mutual compensation between sequence and preexisting chromatin predictors. Ascl1 binding to sites of preexisting active chromatin does not necessarily require strong cognate and secondary sequence motif features, whereas Ascl1 binding to preexisting quiescent or repressed chromatin is correlated with increased motif multiplicity and favorable flanking nucleotide composition.

### Bichrom predicts the relative dependence of neuronal TF binding sites on preexisting chromatin

As demonstrated by our applications to Ascl1 data, Bichrom’s analyses can be interpreted as characterizing a TF’s dependence on preexisting chromatin, both from a global viewpoint and at the level of individual binding sites. We next asked whether we could support such interpretations experimentally. To do so, we apply Bichrom to analyze the binding patterns of TFs that become expressed downstream of Ascl1 in the same mEB-based system^40^.

Specifically, induced Ascl1 (iAscl1) differentiates mEBs into neuronal lineages^40^. Within 12 hours of expression, Ascl1 binding establishes chromatin accessibility at some previously inaccessible binding sites. Ascl1 induces expression of several key neuronal TFs, including Brn2, Ebf2, and Onecut2, within this new chromatin accessibility landscape. Having previously characterized the subsequent genomic binding of Brn2, Ebf2, and Onecut2 using ChIP-seq at iAscl1+48hrs^40^, we applied Bichrom to ask whether each TF’s binding is informed by the chromatin accessibility landscape established by iAscl1 (ATAC-seq, iAscl1+12hrs).

Compared with a network trained using sequence information alone, Bichrom’s incorporation of preexisting iAscl1+12hrs accessibility data significantly improves the representation of iAscl1+48hr binding sites for both Brn2 and Ebf2. Specifically, we observe improvements in recall at a fixed false positive rate (see **Methods, Fig. 6A**) and auPRC (Brn2: sequence network auPRC= 0.23, Bichrom auPRC= 0.35, Ebf2: sequence network auPRC= 0.38, Bichrom auPRC= 0.53; **Supp. Fig. 7**). In contrast, the incorporation of prior chromatin information does not result in improved representation of Onecut2 binding. The sequence-only network and Bichrom show comparable values for both recall (**Fig. 6A**) and auPRC (sequence network auPRC=0.54, Bichrom auPRC=0.56, p-value: 0.59) (**Supp. Fig. 7**). Therefore, Bichrom analysis suggests that Ebf2’s and Brn2’s genomic binding specificity is dependent on preexisting accessibility to a greater degree than that of Onecut2. In other words, Bichrom predicts that Onecut2 has a more pioneer-like behavior than Ebf2 and Brn2.

**Figure 6:**
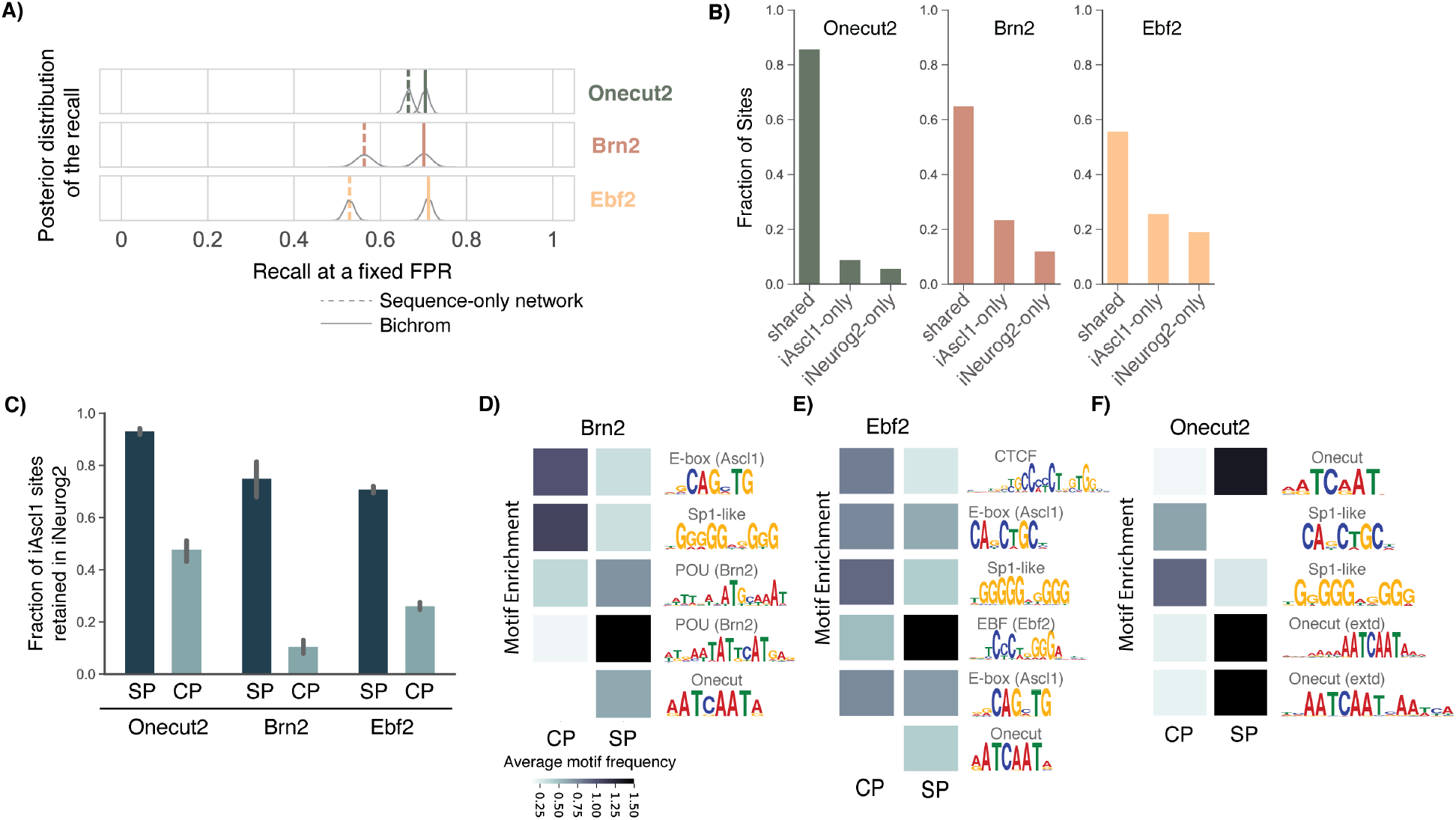
**A)** Bichrom (incorporating preexisting ATAC-seq data) outperforms the sequence-only network at predicting induced Brn2 and Ebf2 binding, but performs at par with the sequence-only network at predicting induced Onecut2 binding. **B)** The percentage of Onecut2, Brn2, and Ebf2 sites that are bound in both iAscl1 and iNeurog2 neurons (shared sites), preferentially bound in iNeurog2 (iNeurog2-only) and preferentially bound in iAscl1 neurons (iAscl1-only). **C)** Bichrom_CHR_ sub-network scores correlate with increasing ATAC-seq tag counts, with scores plateauing for highly accessible regions. MEME-ChIP-determined motif frequency at SP vs. CP sites for **D)** Brn2, **E)** Ebf2, and **F)** Onecut2. For all three TFs, SP sites contain a higher number of cognate motifs. CP and SP sites also display differential secondary motif enrichment.

If Onecut2 is in fact less dependent on preexisting chromatin accessibility, we should expect it to bind many of the same sites in a different chromatin accessibility landscape. Similar to Ascl1, expression of the proneural bHLH TF Neurog2 in mEB cells also leads to differentiation along neuronal lineages, and also causes expression of Brn2, Ebf2, and Onecut2^40^. However, induced Neurog2 (iNeurog2) binds to different sites and thereby establishes a distinct chromatin accessibility landscape (**Supp. Fig. 8**). Consistent with our interpretation of Bichrom’s analysis, the iNeurog2+48hrs binding activities of Brn2 and Ebf2 are affected by the shift in preexisting chromatin accessibility landscapes to a greater degree than Onecut2. Specifically, only 65% of Brn2 sites and 56% of Ebf2 sites are shared between iAscl1 and iNeurog2 cell types, whereas 86% of Onecut2 binding sites are shared (**Fig. 6B**)^40^. This comparative analysis of differential binding across two cell types thus supports Bichrom’s ranking of the TFs’ dependence on preexisting chromatin, which was derived from analysis of ChIP-seq data from a single cell type (i.e., iAscl1+48hrs). Furthermore, Bichrom’s prediction that Onecut2 is less dependent on preexisting chromatin is consistent with recent literature that establish Onecut TFs as neuronal pioneers^51,52^.

Turning to the predictors of individual TF binding sites, Bichrom’s two-dimensional latent space embeddings again show wide ranges of Bichrom_SEQ_ and Bichrom_CHR_ sub-network scores for all three neuronal TFs (**Supp. Fig. 9**). For each TF, we extract subsets of binding sites that we define as “sequence-predicted” (SP; sites in top 25^th^ percentile of Bichrom_SEQ_ scores and bottom 25^th^ percentile of Bichrom_CHR_ scores) and “chromatin-predicted” (CP; sites in top 25^th^ percentile of Bichrom_CHR_ scores and bottom 25^th^ percentile of Bichrom_SEQ_ scores). If our latent space embeddings are meaningful, we would expect the “CP” sites to be more dependent on a shift in the preexisting chromatin landscape than the “SP” sites. Analysis of differential binding downstream of iNeurog2 again supports our interpretation. For all three neuronal TFs, “SP” iAscl1 binding sites show consistently higher levels of retention in iNeurog2 cells than “CP” sites (**Fig. 6C**). For example, 74% of Brn2’s iAscl1 SP sites are also bound in iNeurog2 cells, while only 11% of Brn2’s iAscl1 CP sites are retained (**Fig. 6C**). Thus, Bichrom’s prediction that some individual binding sites are more dependent on preexisting chromatin features is confirmed by those sites being more sensitive to a shift in the underlying chromatin landscape.

Our interpretations of the latent embeddings are also supported by motif analysis at SP and CP sites (**Fig. 6D-F**). The cognate DNA-binding motifs for Brn2, Ebf2, and Onecut2 show consistently higher enrichment in the SP sites compared with their CP sites (**Fig. 6D-F**). In contrast, and even though the Bichrom_CHR_ subnetwork does not use any DNA sequence information, we find that all three TF’s CP sites contain higher enrichment for a Sp1-like motif and the CAGSTG E-box motif that is preferred by Ascl1. The latter observation suggests that some CP sites bound by the neuronal TFs at 48hrs may be made accessible by Ascl1 binding at 12hrs. Indeed, we find that CP sites overlap preexisting Ascl1 sites (iAscl1+12hrs) at a significantly higher rate than SP sites: for example, 59% of Brn2’s CP sites overlap preexisting Ascl1 binding compared to 1% of Brn2 SP sites. Similarly, we find a CTCF-like motif enriched at higher rates in Ebf2 CP sites than SP sites (**Fig. 6E**), suggesting that CTCF binding in a prior cell stage may establish a favorable chromatin environment for Ebf2 binding. Comparing with mES CTCF ChIP-seq data, we find that 33% of Ebf2 CP sites overlapped with CTCF binding events, while fewer than 1% of Ebf2 SP sites do so.

In summary, the latent network embeddings identify sites that are more likely to be differentially bound in distinct chromatin environments. Additionally, they can be used to identify the diverse sequence, chromatin, and co-factor feature compositions that specify genome-wide TF binding.

### The predictive capacity of preexisting chromatin varies across TFs

Finally, we asked whether the ability of preexisting chromatin to explain binding specificity varies across a broader range of TFs, cellular conditions, and datasets. We applied Bichrom to analyze the binding of 12 induced TFs from studies where aspects of the preexisting chromatin environment were also characterized. Specifically, we used Bichrom to analyze the binding of 9 TFs that were induced in mouse NIH-3T3 fibroblasts and assayed approximately 12 hours post TF induction (**Supp. Table 3**)^45^, using NIH-3T3 ATAC-seq to define the preexisting chromatin accessibility landscape (**Supp. Table 4**). We also analyzed the binding of three previously established human pioneer TFs – OCT4, GATA4, and FOXA2 - that were induced in human BJ fibroblasts and assayed 4 days post induction^53^. In these latter analyses, the preexisting chromatin landscape was defined by ATAC-seq, H3K27ac, H3K4me2, and H3K27me3 (**Supp. Table 5**).

Applied to the NIH-3T3-induced TFs, Bichrom’s incorporation of preexisting accessibility increased predictive accuracy for NIH-3T3 induced Bhlhb8, Sox15, Sox2, Dlx6, Rhox11, Hlf, and FoxA1 (**Fig. 7A, Supp. Fig. 10A**). However, the incorporation of preexisting accessibility did not significantly increase predictive accuracy for Duxbl or Cdx2, suggesting that the binding patterns of these TFs are not as dependent on the preexisting chromatin landscape (**Fig. 7A, Supp. Fig. 10A**). In the human fibroblast datasets, we find the preexisting chromatin data improves Bichrom’s predictions of induced TF binding to a limited degree for FOXA1 and GATA4, but does not result in improved predictive capacity for OCT4 (**Fig. 7B, Supp. Fig. 10B**). Therefore, our computational framework suggests that the preexisting chromatin environment can predict the binding of TFs to varying degrees.

**Figure 7:**
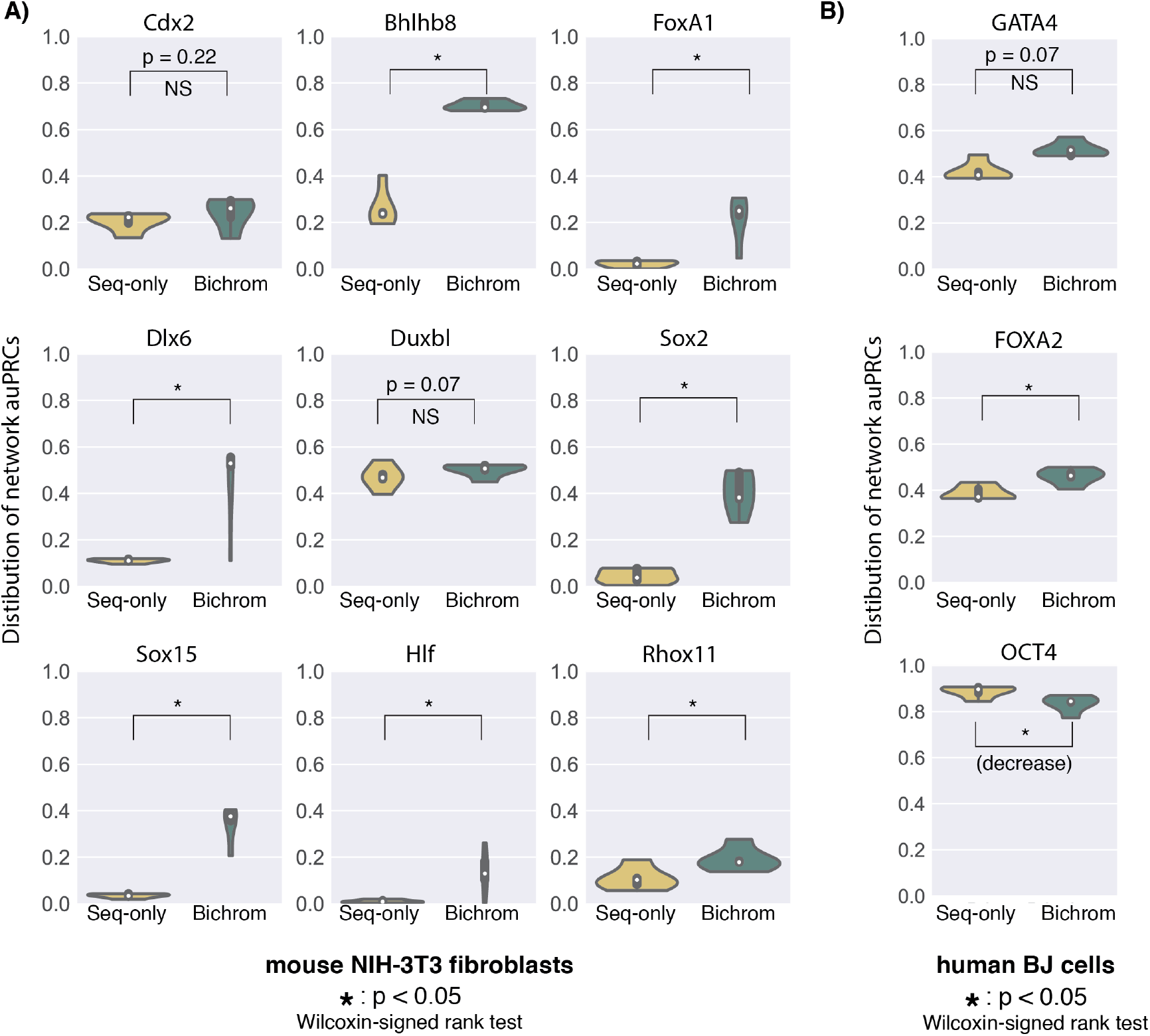
**A)** Distribution of model performance (auPRC) for a sequence-only network and Bichrom (sequence and preexisting NIH-3T3 ATAC-seq) for 9 TFs induced in mouse NIH-3T3 fibroblasts. The boxplots represent data from 5 independent training sets; each consisting of a separate held-out test chromosome. **B)** Distribution of model performance (auPRC) for a sequence-only network and Bichrom (sequence and preexisting fibroblast ATAC-seq, H3K27ac, H3K4me2, and H3K27me3) for FOXA2, GATA4, and OCT4 induced in human BJ fibroblasts.

Using Bichrom’s latent embeddings of individual binding sites, we interpret the sequence and preexisting chromatin features that predict DNA binding of the examined induced TFs. We find that induced TF binding sites are assigned a broad range of Bichrom_SEQ_ sub-network scores (**Supp. Fig. 11A**). For the majority of TFs, sites scored highly by the Bichrom_SEQ_ subnetwork are associated with increased cognate motif enrichment (**Supp. Fig. 12**). For the TFs FoxA1 and Duxbl, higher scoring sites are associated with an increased cognate motif enrichment as well as cognate motifs that contain additional informative flanking nucleotide sequences (**Supp. Fig. 12**).

Furthermore, for a majority of TFs that bind at least 10% of their sites in previously inaccessible chromatin (Duxbl, Cdx2, Bhlhb8, FOXA2, GATA4), we find that sites bound in pre-inaccessible chromatin are on average assigned higher Bichrom_SEQ_ scores, suggesting that there exists some degree of compensation for these TFs where pre-accessible chromatin enables binding at sites with weaker sequence features (**Supp. Fig. 11B**). The only TF for which this pattern is reversed is OCT4. Instead of the canonical OCT4 motif, Bichrom_SEQ_ scores are associated with GC-rich sequences (**Supp. Fig. 12**). In this case, previously accessible sites are assigned higher Bichrom_SEQ_ scores (associated with GC-rich sequences) than previously inaccessible sites, which are enriched for a SOX-related motif (**Supp. Fig. 11B, 12**). These results are consistent with the observation from Donaghey *et al*. that 73% of OCT4 ChIP-seq peaks in their human fibroblast inductions overlap CpG islands^53^.

Taken together, our analyses of a broad range of induced TFs demonstrate that Bichrom is a useful tool for analyzing the global preexisting chromatin predictors of induced TF binding, and for examining the diversity in sequence and preexisting chromatin features that define individual TF binding sites.

## Discussion

TFs bind subsets of their cognate motif instances in a cell type-specific fashion. Such specificity in TF binding results from an interplay between the TF’s inherent sequence preferences and cell type-specific chromatin landscapes^6,12^. The question naturally arises as to which local chromatin features might enable or inhibit a given TF’s DNA binding activities. However, if we can measure a TF’s binding occupancy using ChIP-seq, it has by definition already had its own impact on chromatin in that cell type (e.g. by making its binding sites accessible or by recruiting histone modification enzymes). Concurrent chromatin landscapes therefore predict TF binding in the same cell type^54,55^, but cannot be used to model the causal determinants of that binding.

Bichrom represents an interpretable neural network architecture that can be used to assess the relative contributions of DNA sequence and preexisting chromatin features in specifying an induced TF’s genomewide binding sites. Related to our work, several previous studies have assessed the effects of prior chromatin landscapes on the binding of specific TFs^11,12,44,53,56,57^. Our work aims to provide a unified predictive framework for quantifying and formalizing the relative contributions of sequence and chromatin predeterminants to TF binding across a range of TFs.

We have demonstrated that comparing Bichrom’s TF binding predictive performance to that of a sequence-only neural network allows us to infer a TF’s overall dependence on preexisting chromatin environments. While incorporating prior chromatin data significantly improves predictive performance for some TFs (e.g., Brn2, Dlx6, and Bhlhb8), others seem to display less dependence on prior chromatin (e.g., Onecut2, Duxbl, and Cdx2). Thus, our approach may offer a metric for quantifying the “pioneering” activity of a TF; those that display less dependence on preexisting chromatin may be more likely to override those landscapes. We allow that our estimates of a TF’s dependence on preexisting chromatin may still be cell typespecific as opposed to an innate feature of a given TF. For example, while our analyses suggest that binding sites for the well-characterized pioneer factor FoxA1 are dependent on prior chromatin, this may be specific to the measured context of NIH-3T3 cells. It is possible, for instance, that TFs that cooperate with or otherwise predict FoxA1 binding are already present in NIH-3T3 cells, and FoxA1 may be less dependent on preexisting chromatin in other cell types.

We further demonstrate the ability of the bimodal network to deconvolve the sequence and prior chromatin features that predict TF binding at individual sites. Interpretation of Bichrom network embeddings at individual binding sites suggests that sequence and preexisting chromatin landscapes are not independent predictors of TF binding. Rather, sequence and preexisting chromatin are mutually compensatory features that define a continuum of sites that may be bound by the induced TF. While genomic loci with weaker sequence signatures may be bound by TFs given a favorable local chromatin environment, the same signatures might not be sufficient to drive TF binding at inaccessible or unfavorable chromatin. For example, Ascl1 is more likely to bind pre-inaccessible loci in the presence of certain sequence features such as high motif multiplicity and favorable motif flanks.

On the other hand, some highly accessible active promoters and enhancers are bound even with weaker sequence signatures, as defined by low activation scores from the Bichrom_SEQ_ sub-network in our model. We note that some TF-bound regions with high preexisting chromatin (Bichrom_CHR_) sub-network activations and low sequence (Bichrom_SEQ_) sub-network activations might represent artifactual ChIP-seq enrichment^59^. Alternatively, these regions may represent direct binding to weaker motifs, or indirect binding mediated by interactions with mES or NIH-3T3 cell regulators^19^. While previous studies have proposed sequence-conditional binding to inaccessible chromatin for a few TFs^12,39,60^, our work suggests that this compensatory mechanism may exist across a broader range of TFs.

Finally, our results quantify the degree to which different TFs may have different abilities to interact with preexisting chromatin landscapes^42,61,62^. It will therefore be of interest to examine how the relative contributions of sequence and preexisting chromatin vary in determining the binding of a wider range of TFs, and across a wider array of cell types. Identifying such sequence and chromatin predeterminants of TF binding will be crucial for understanding gene regulation in various dynamic systems such as development and cellular programming.

## Methods

### ChIP-seq & ATAC-seq data (mEB cells)

Generation of the inducible iAscl1 and iNeurog2 mouse ES cell lines and corresponding ChIP-seq data is more completely described in Aydin, *et al*.^40^. Briefly, inducible cell lines were generated using the inducible cassette exchange (ICE) method as previously described^63^. TF gene constructs are inserted in single copy into the expression-competent HPRT locus. The resulting iAscl1 and iNeurog2 ES cells are differentiated on untreated plates for 2 days to form embryoid bodies, and then expression of the transgene is induced via Doxycycline. Ascl1 and Neurog2 binding was assayed by ChIP-seq 12 hours after Dox induction using the anti-Ascl1 (Abcam, ab74065) and anti-Neurog2 (Santa Cruz, SC-19233) antibodies. We assayed histone modifications as well as chromatin accessibility in EBs with ChIP-seq and ATAC-seq, respectively (Suppl. Data Table 1). We collected additional publicly available histone modification and histone variant ChIP-seq datasets from mouse ES cells (Suppl. Data Table 2). Together, our dataset defining the chromatin environment of mouse pluripotent cells consists of the following 12 data types: ATAC-seq, H2A.Z, acH2A.Z, H3K27ac, H3K27me3, H3K9me3, H3K4me1, H3K4me2, H3K4me3, H3K9ac, H4K20me3 and H3K36me3.

### ChIP-seq & ATAC-seq data (NIH-3T3 cells and BJ cells)

ChIP-seq data for TF inductions in mouse NIH-3T3 fibroblasts was retrieved from Raccaud *et al*. (GSE119784)^45^. We filtered for TFs that were not expressed as defined by RNA-seq in the NIH-3T3 cell line^45^. We used NCIS to estimate the sequenced control-based normalization factors for each TF ChIP-seq experiment^64^. Further, we filtered out induced TFs that had a MultiGPS-reported signal fraction < 0.01 and were single-replicate ChIP-seq experiments (Suppl. Data Table 3). We used five ATAC-seq experiments (Suppl. Data Table 4) as replicates to construct the network’s ATAC-seq input^45^. ChIP-seq data for FOXA2, GATA4, and OCT4 inductions in human BJ fibroblasts was retrieved from Donaghey *et al*. (GSE110214)^23^. ATAC-seq, H3K27me3, H3K4me2 and H3K27ac were used to construct the network’s chromatin input.

### TF ChIP-seq & preexisting chromatin data processing

Fastq files were aligned to the mouse (version mm10) or human (version hg38) genomes as appropriate using Bowtie (version 1.0.1)^66^ with options “-q -best --strata -m 1 --chunkmbs 1024”. Only uniquely mapped reads were considered for further analysis. MultiGPS^65^ (version 0.74) was used to define transcription factor DNA binding events, and was run with default options except for “ --fixedpb 5” and excluding ENCODE blacklist regions. A q-value cutoff of 0.01 (assessed using binomial tests and Benjamini-Hochberg multiple hypothesis test correction) was used to call statistically significant binding events with respect to sequenced input material collected from the same cell line. Peak-finding statistics are reported in Suppl. Table 3. Paired-end ATAC-seq reads were aligned using Bowtie2 (version 2.2.2) using the “-q --very-sensitive” options^67^. ChromHMM^50^ (version 1.2.0) was run using default parameters.

### Training and test set construction

For testing, we divided the genome into 500bp non-overlapping windows. For training, we use 500bp overlapping windows, each of which are sequentially offset by 50bp. Genomic windows overlapping peak calls with a *p*-value ≤ 0.001 are labeled as bound. Windows overlapping non-significant peaks from MultiGPS are labeled ambiguous. All other genomic windows (~99%) are labeled as unbound. The sequence sub-network (Bichrom_SEQ_) takes as input 500bp sequences. Each nucleotide is encoded as a one-hot vector, such that only the index corresponding to the input nucleotides is set to one, and all other indices are set to zero. For each chromatin input data track, we extract the per-base read counts at each genomic locus. These raw coverage counts are binned into ten 50bp non-overlapping bins (covering 500bp windows). The binned read counts are total tag normalized for each replicate, and we use the replicate average at each bin as input to our network. The chromatin datasets are stacked, resulting in a 10 x *k* chromatin input, where *k* is the number of assayed histone modifications/chromatin accessibility. For analyses using only prior ATAC-seq (i.e., iAscl1+12hr, iNeurog2+12hr, and NIH-3T3 induced TFs): *k*=1. For analyses of the mEB-induced TFs using prior ATAC-seq and other histone modification data: *k*=12. For analyses of human BJ fibroblast-induced TFs: *k*=4.

### Neural network architecture

The bimodal network architecture was designed in order to produce interpretable latent embeddings. Since hybrid CNN-LSTM networks have previously been shown to perform well at TF binding prediction tasks, we chose to use a CNN-LSTM based architectures for Bichrom_SEQ_ and Bichrom_CHR_^24,46^. The parameters of the individual sub-networks (Bichrom_SEQ_ and Bichrom_CHR_) were selected using a limited hyper-parameter random grid search (chr10 held-out test set, chr17 validation). We used a random grid-search over the (1) number of dense layers, (2) size of the dense layers, (3) number of convolutional filters, (4) activation functions and the (5) dropout rate (see tested parameter values in **Supp. Table 6**). To test whether our selected CNN-LSTM based architecture performed at par with alternative CNN-based architectures (e.g., deeper multi-layer CNNs), we generated a range of network architectures by randomly selecting hyper-parameter value combinations over (1) number of convolution layers, (2) convolutional kernel size, (3) number of convolutional filters, (4) max pooling size, (5) max pooling stride, (6) number of dense layers, (7) number of dense nodes and (8) dropout rate to select network hyper-parameters (see tested parameter values in **Supp. Table 7**). We found that Bichrom’s CNN-LSTM network performed comparably to architectures with multiple convolutional layers when predicting Ascl1 binding in mEB cells (**Supp. Fig. 1**).

In the Bichrom_SEQ_ sequence sub-network, the 500bp, one-hot encoded sequence input is first subjected to a 1-dimensional convolution layer, with each index in the one-hot encoding acting as a channel into this convolution. The convolutional layer consists of 240 x 20bp long filters. The convolutional filters within 15bp intervals are max-pooled, and the pooled convolutional output is used as input into a long short-term memory (LSTM) layer. The LSTM outputs a 32-vector, which then passes through two dense layers, both subjected to ReLU activation and dropout. The activations from the final dense layer are input into a single tanh activated dense node. The Bichrom_CHR_ chromatin sub-network uses convolutional filters that span two input bins. The filters are followed by an LSTM to model any observable tag densities discriminative of TF binding. The LSTM activations are input into a single dense layer followed by a single tanh activated dense node. The activations of both sub-networks are weighted by a final sigmoid activated node, used to output binding probability. The network is trained to predict ChIP-seq by minimizing the binary cross-entropy loss *J*:

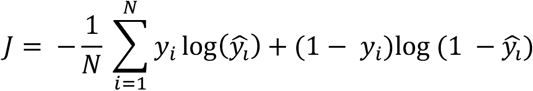

Area under the precision-recall curve is used as a metric to measure network performance. Chromosome 17 is held out as a validation chromosome. Chromosomes 10-16 and chromosomes 18-19 are sequentially held out as test chromosomes in a *k*-fold training procedure.

### Neural network training strategies

To prevent the sequence-only network from learning accessibility-related sequences, we customize the sampling used to construct mini-batches for gradient descent-based training. We construct training batches such that in each batch, either only accessible or only inaccessible bound and unbound training data is used. This sampling strategy reduces model false positives at preexisting accessible regions, and leads to an improvement in sequence-only model performance measured on held-out chromosomes, as measured via the area under the precision recall curve (auPRC) (**Suppl. Fig. 4A, B**). For TFs with less than 10% of their binding sites in previously inaccessible chromatin, a chromatin-matched training results in poor predictive performance due to the small number of positive training examples in previously inaccessible chromatin. For these TFs, we use a less stringent strategy wherein within each batch, the fraction of previously accessible sites is similar in both the bound and unbound training instances.

The bimodal network aims to learn both sequence and prior chromatin signatures that characterize genome-wide TF binding. We can thus no longer control for accessibility distributions across bound and unbound training sets to prevent spurious learning of prior chromatin-related sequence signatures when training the bimodal network. To address this problem, we transfer weights from the previously trained sequence-only network to the Bichrom_SEQ_ sequence sub-network in Bichrom’s bimodal network. While the lower-level layer sequence sub-network weights are kept fixed during Bichrom training, the weights for the final dense layer in the Bichrom_SEQ_ sub-network are allowed to vary to fit the genome-wide TF binding data.

Keeping the convolutional kernels fixed while re-training the final dense layers should allow the network to optimally predict TF binding, without learning new accessibility-related sequence features^68^. However, to ensure that the compensatory sequence-chromatin behavior observed across TFs is not due to such a network parameterization, we train a control network in which all dense weights within the Bichrom_SEQ_ subnetwork are kept fixed. We find that the 2-dimensional embeddings for each TF binding site retain the same compensatory pattern as observed earlier, suggesting that network parameterization is not responsible for the observed sequence-chromatin compensation (**Supp. Fig. 5**). Bichrom’s joint bimodal network with fixed convolutional kernels can therefore be trained using imbalanced batches constructed by random sampling unbound data across the genome.

### Benchmarking against the ENCODE DREAM challenge data

The ENCODE DREAM challenge data was downloaded from https://www.synapse.org/#!Synapse:syn6131484/wiki/402026. The chromosomes available for training by the challenge were chromosomes chr2-7, chromosomes 9-20 and chromosomes 22 and X. Methods described in the ENCODE-DREAM challenge were tested on chromosomes 1, 8 and 21. As the challenge held-out data is not been made available at time of writing, we additionally held-out chromosome 18 from our training set in order to test our models. We used DNA sequence and concurrent DNase-seq as inputs into the bimodal network. We used the window definitions derived from the challenge (200 base pair windows, 50 base pair step size) in order to be consistent with other methods in the challenge. Notably, we did not make use of concurrent RNA-seq data or *in vitro* PBM-derived TF motif data that was used by several other methods. We evaluated network performance using the auPRC, and compared our results to those reported for the top-performing methods in the challenge (**Supp. Fig. 2**).

### Deriving the latent network embeddings

Let *ϕ_S_* and *ϕ_C_* represent the non-linear transformations applied to sequence feature vectors ***X_S_*** and chromatin feature vectors ***X_C_*** respectively. Then, for each input genomic window *i*, 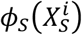 and 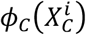 represent real-valued outputs of the sequence and chromatin sub-networks at that window.

The bimodal network models the network output ***y*** as follows:

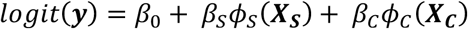

Here, *β_S_* and *β_C_* are weights assigned to sequence and chromatin sub-networks respectively, and *β*_0_ is the bias term. Due to the linear relationship between *logit*(***y***) and the transformed predictors, the weighted subnetwork activations can be interpreted as the relative contributions of each modality to the networks output probability at a window *i*:

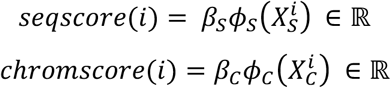

Therefore, the network embeds each input data point *i* in a 2-dimensional space defined by *seqscore*(*i*) and *chromscore*(*i*).

### Feature attribution with integrated gradients

We use integrated gradients^48^ (IG) to estimate the relative importance of each nucleotide 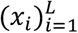 within each input sequence ***x*** of length *L* base pairs. Due to the nonlinearity in neural networks, it is not trivial to assign relative feature importance using the feature-associated network weights. Instead, gradient-based techniques are often used in order to estimate relative feature importance. Vanilla gradients estimate feature importance as the partial derivative of the network output with respect to each input feature (in our case nucleotide *x_i_*). However, due to the presence of multiple saturating functions used in neural networks, vanilla gradients do not satisfy the property of sensitivity. In other words, they are not guaranteed to assign non-zero attribution to all features *x_i_* that alter the network output when compared to reference feature vectors that inherently produce zero output probabilities. Integrated-gradients considers how predictions at input feature vectors differ from reference feature vectors. More specifically, integrated gradients calculate the gradients at all points along a straight-line path from the reference feature vector ***x^b^*** to the input feature. In our case, we define a reference feature as a sequence vector such that at each position, each nucleotide is equally likely. In other words, our reference sequence is a 4 * 500 matrix, with each column defined as [0.25, 0.25, 0.25, 0.25]. We implemented integrated gradients as defined in Sundarajan *et al*.^48^

### Analysis of sequence and preexisting chromatin predictors of Ascl1 binding sites

As described above, we applied integrated gradients (IG) to identify the sequence motifs driving sequence sub-network scores for Ascl1^48^. However, attribution methods typically operate on individual sequences. To overcome the local behavior of such attribution methods, we extracted 20 bp sequences surrounding the local IG peaks; i.e., we extracted regions or “hills” that drive the neural network output from each sequence bound by Ascl1. We clustered the IG-derived hills based on their underlying sequence features using a K-means based clustering procedure described previously in SeqUnwinder^69^. Briefly, only *k*-mers present in at least 5% of IG-derived hills are used for clustering, and K-means clustering is performed using Euclidean distance as a metric^69^. We then perform motif discovery using MEME^70^ to identify the enriched motifs within each K-means defined cluster^69^.

To investigate the correlation between Bichrom_SEQ_ scores and Ascl1 cognate motif multiplicity, we divided bound loci into categories based on their motif multiplicity, and measured the Bichrom_SEQ_ subnetwork scores for each category. Only 5% of all Ascl1-bound sites lacked exact matches to the core Ascl1 Ebox motif CAGSTG, and these sites were assigned the lowest median scores by the sequence sub-network (**Fig. 4C**). To ensure that increased sequence network scores at Ascl1 binding sites with multiple motifs do not stem from the presence of confounding features at these sites, we directly tested the relationship between motif multiplicity and Bichrom_SEQ_ scores. We inserted between one and five randomly spaced CAGSTG *k*-mers in a set of 10,000 randomly generated 500 bp sequences (background frequencies A/T=0.5 and G/C=0.5). We divided the simulated 500 bp sequences into categories based on the number of embedded CAGSTG *k*-mers. Each simulated sequence was input into a Bichrom network trained using Ascl1 ChIP-seq data and the distribution of Bichrom_SEQ_ sub-network scores at each sequence category was calculated (**Fig. 4D**).

To investigate the effect of various nucleotides flanking the Ascl1 cognate motif, we constructed a sequence in which each position is encoded as a [0.25,0.25,0.25,0.25]^T^ vector; i.e. each base [A, T, G, C]^T^ occurs with equal probability at each position in this sequence. We inserted CAGSTG *k*-mers flanked by variable nucleotides into this reference sequence, and again scored them with the Ascl1-trained Bichrom_SEQ_ subnetwork.

To investigate the relationships between Bichrom_CHR_ scores at Ascl1 binding sites and individual preexisting chromatin features (**Fig. 5A**), we divided Ascl1 binding sites into quartiles based on their Bichrom_SEQ_ scores and calculated the mean normalized tag enrichment for each preexisting chromatin feature at each quartile. Further, we identified genome-wide enrichment domains were using the DomainFinder utility in SeqCode (https://github.com/seqcode/seqcode-core/blob/master/src/org/seqcode/projects/seed/DomainFinder.java) and calculated Bichrom_CHR_ scoring distributions at all preexisting chromatin domains for each input chromatin dataset (**Fig. 5B**). We also calculated the distribution of Bichrom_CHR_ and Bichrom_CHR_ scores Ascl1 binding sites categorized into 12 states based on ChromHMM state calls (**Fig. 5C**).

### Motif Discovery

*De novo* motif discovery for iAscl1 induced Ebf2, Onecut2, and Brn2, the mouse NIH-3T3 induced TFs and the human BJ induced TFs was performed using MEME-ChIP^71^ (version 5.1.0) with default settings. All motifs with MEME E-values < 0.01 discovered at SP and CP sites were reported. Repetitive polyA or poly-T repeats were excluded. Motif scanning was performed using FIMO^72^, all hits greater than the default p-value threshold of 1e-4 were reported.

### The posterior distribution of the model recall

We used the model recall at a fixed false positive rate (FPR) to compare model performance across TFs. TPs are true positives in the held-out test set, whereas FNs are false negatives in the test set.

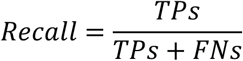

However, we note that ChIP-seq signal fractions and the number of peaks called vary widely across TFs. Models trained to predict binding for TF ChIP-seq experiments that contain smaller numbers of peaks (and correlated lower signal fractions) suffer from having access to limited training data. In order to quantify our confidence in the model recall, we use a probabilistic framework that models the recall for each TF given the underlying ChIP-seq data. Specifically, analogous to Brodersen *et al*.^73^, we consider the observed model recall (measured on a single held-out test chromosome) to be an actualization of an underlying true recall value *r* given *N* independent Bernoulli trials, where *N* is the number of binding sites in the held-out test chromosome. Each binding site can be either labeled a true positive (success) or a false negative (failure) by the network.

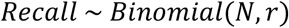

We derive the posterior distribution of the recall *r* assuming a beta prior (parameters *a*=1, *b*=1; equivalent to a uniform prior - for details, see Brodersen *et al*.^73^). The mode of this posterior distribution is the observed model recall. If a TF ChIP-seq experiment contains a small number of peaks, the distribution of *r* has high variance (e.g. Brn2, **Fig. 6A**). On the other hand, a low variance in the distribution of *r* reflects a higher degree of confidence in our estimate of the recall (e.g. Ebf2, Onecut2, **Fig. 6A**).

### Availability

Open source code (MIT license) is available from https://github.com/seqcode/iTF. ChIP-seq data have been uploaded to GEO under accession GSE114176.

## Acknowledgements

This manuscript is based upon work supported by the National Science Foundation ABI Innovation Grant No. DBI1564466 (to SM). Any opinions, findings and conclusions or recommendations expressed in this material are those of the authors and do not necessarily reflect the views of the NSF. This work was also supported by the Academic Computing Fellowship (to DS), NIGMS R01GM121613 (to SM), NICHD R01HD079682 (to EOM), and an NVIDIA GPU equipment grant. We thank Chunyu Ma for a critical reading of the manuscript.

## Supplementary Material

**Supplementary Figure 1:**
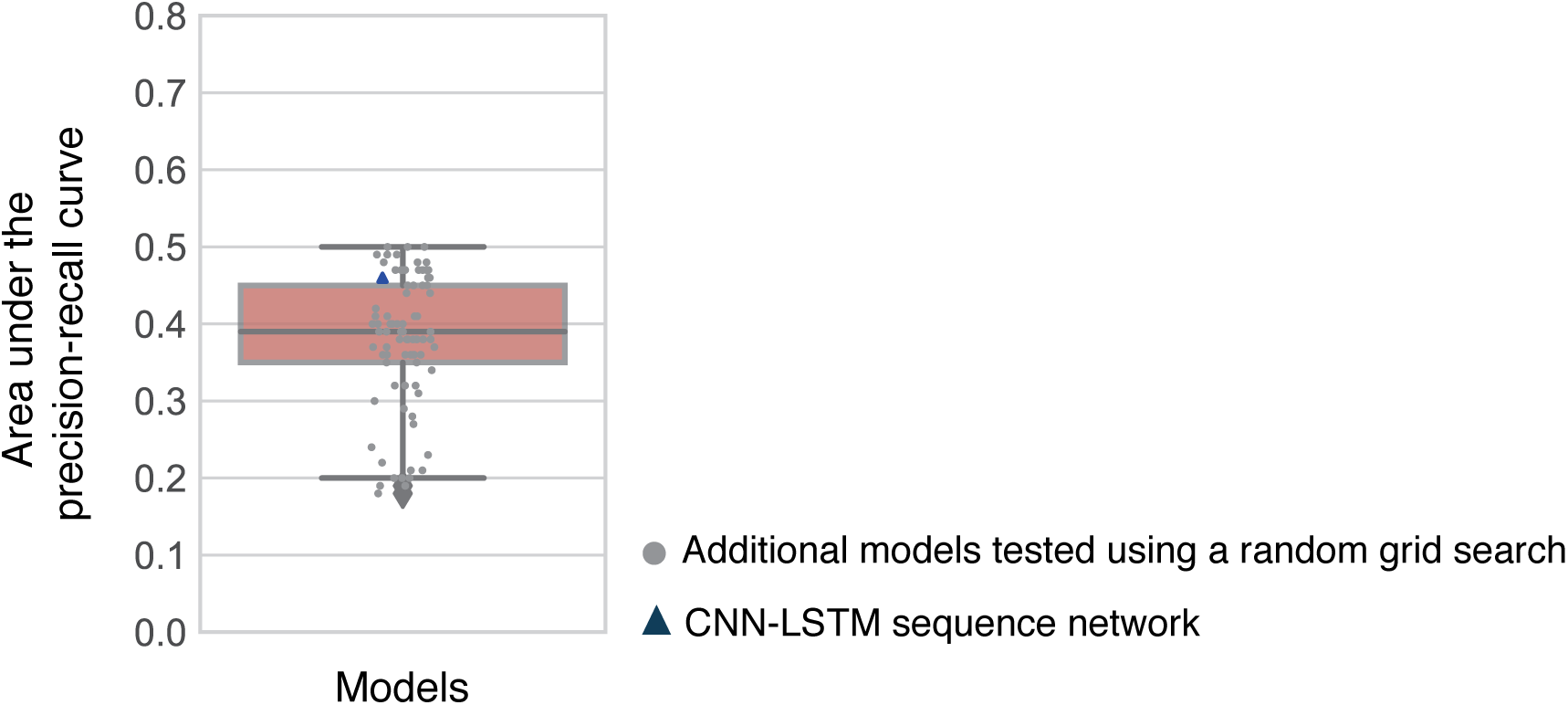
Comparison of performance between the bimodal CNN-LSTM architecture employed by Bichrom and bimodal CNN architectures with varying hyper-parameters and numbers of layers. Performance is assessed as area under the precision-recall curve when predicting Ascl1 binding (mEB+12hrs) using sequence and preexisting mEB and mES chromatin features (chr10 held-out test set, chr17 validation). The alternate CNN architectures were chosen using a random grid search over the following hyper-parameters: (1) number of convolution layers, (2) convolutional kernel size, (3) number of convolutional filters, (4) max pooling size, (5) max pooling stride, (6) number of dense layers, (7) number of dense nodes and (8) dropout rate to select network hyper-parameters (search values shown in Supp. Table 7).

**Supplementary Figure 2:**
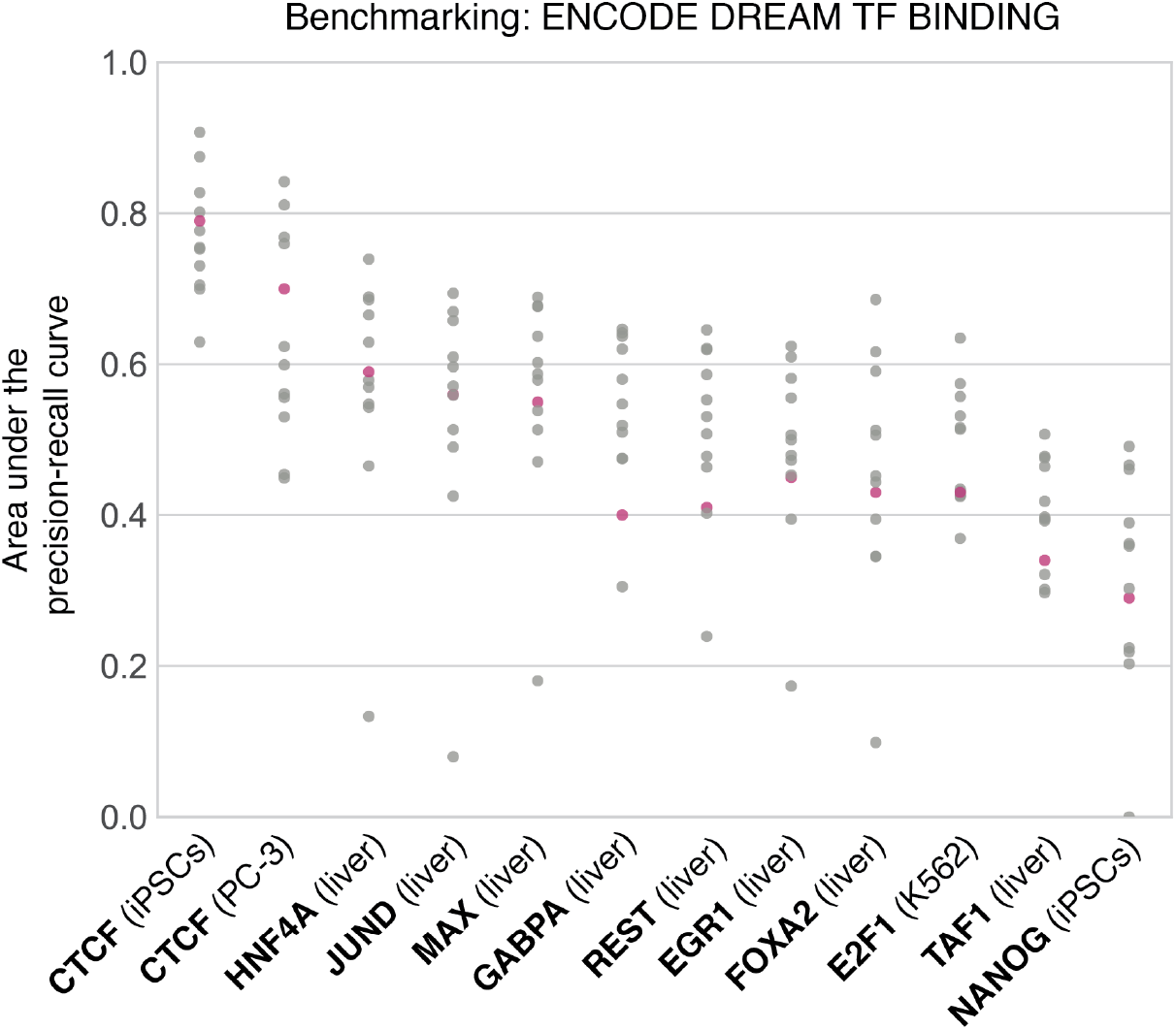
Area under the precision-recall curve for the Bichrom architecture (red) run using sequence and concurrent chromatin accessibility (DNase-seq) as inputs for predicting within-cell-type genome-wide TF binding, compared to auPRCs of the top 10 methods from the ENCODE-DREAM TF binding prediction challenge.

**Supplementary Figure 3:**
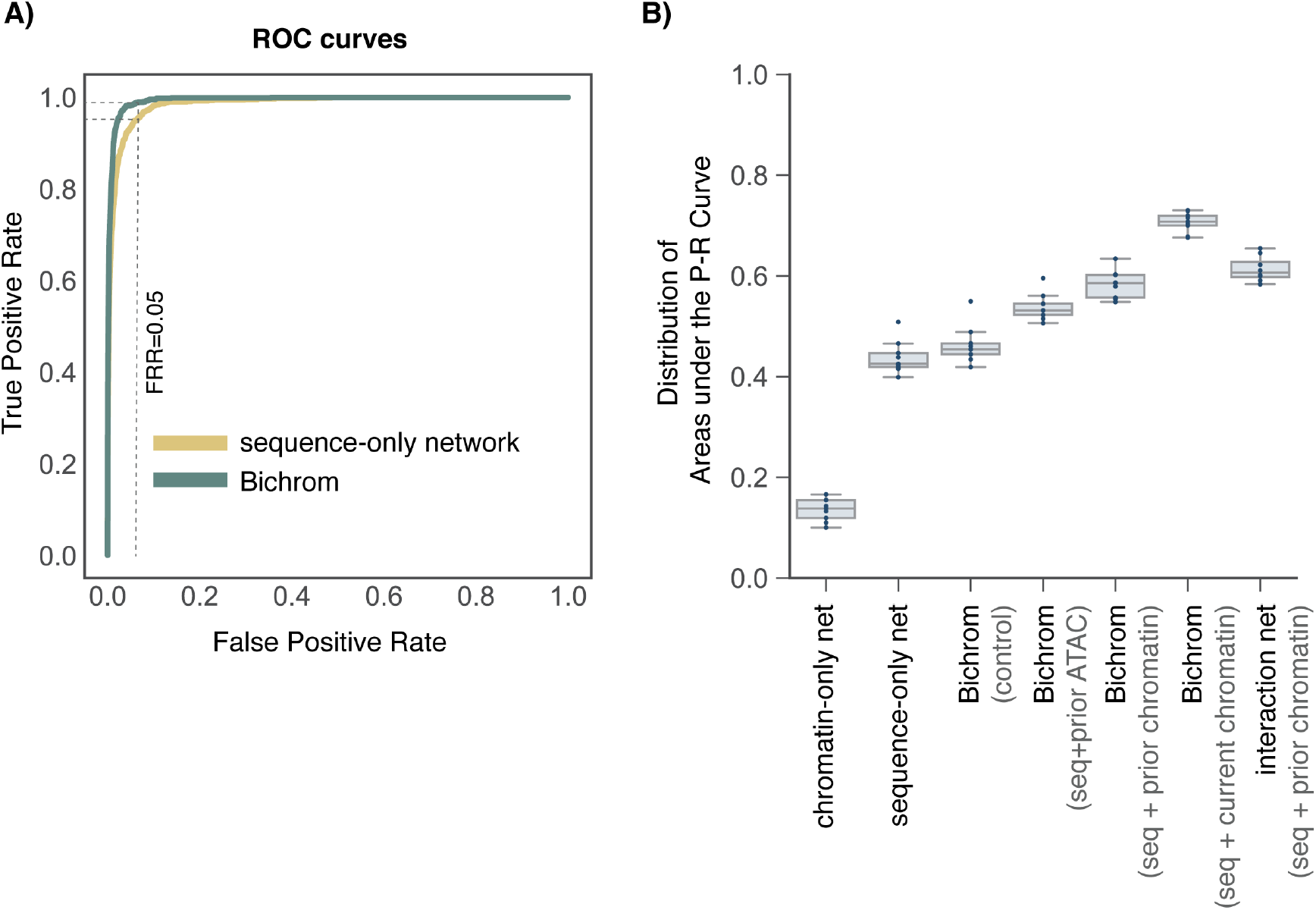
**A)** The receiver-operating characteristic (ROC) curves for the sequence-only network and Bichrom trained to predict Ascl1 binding. At a fixed false positive rate (FPR) of 0.05, both models have high true positive rates (TPR > 0.9). **B)** Model auPRC distributions for various neural networks over 9 independent training sets. The performance of Bichrom’s additive bimodal network architecture is comparable to that of an early integration network (interaction net), confirming that an additive model does not result in a loss of predictive accuracy.

**Supplementary Figure 4:**
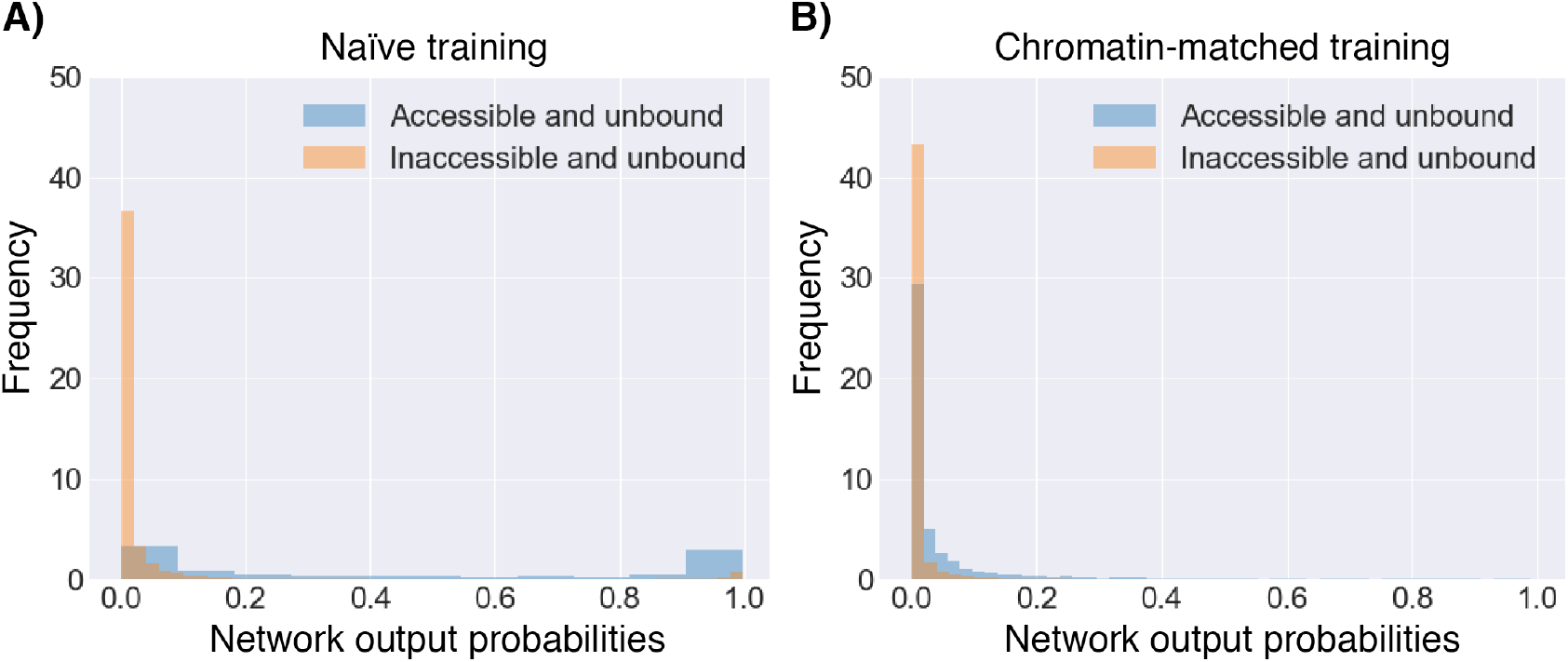
The distribution of Bichrom_SEQ_ sub-network scores at Ascl1-unbound sites divided by prior accessibility. **A)** In the naïve training, a large number of pre-accessible unbound regions (blue) are incorrectly assigned high Bichrom_SEQ_ scores. **B)** This bias is lost in the chromatin-matched training approach, with the model behaving more uniformly across pre-accessible and pre-inaccessible sites.

**Supplementary Figure 5:**
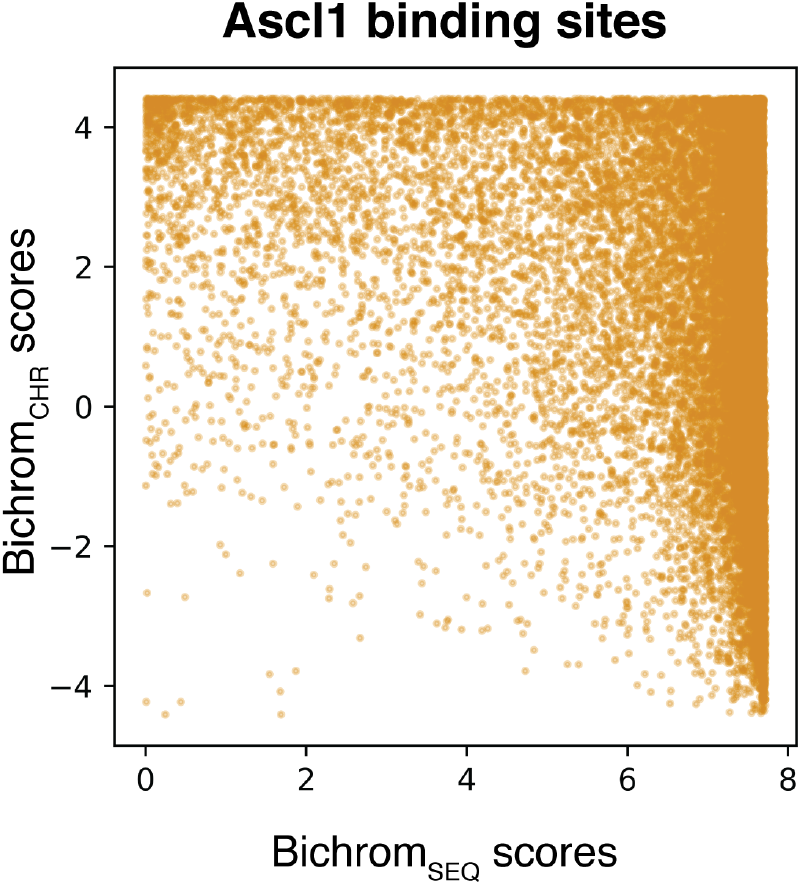
Sequence and prior chromatin sub-network embeddings for Ascl1 from a control bimodal neural network in which all sequence sub-network weights are kept fixed (non-trainable) while training the network on genome-wide binding data (bound genomic sites as positive training examples and a larger set of randomly selected unbound sites are negative training examples).

**Supplementary Figure 6:**
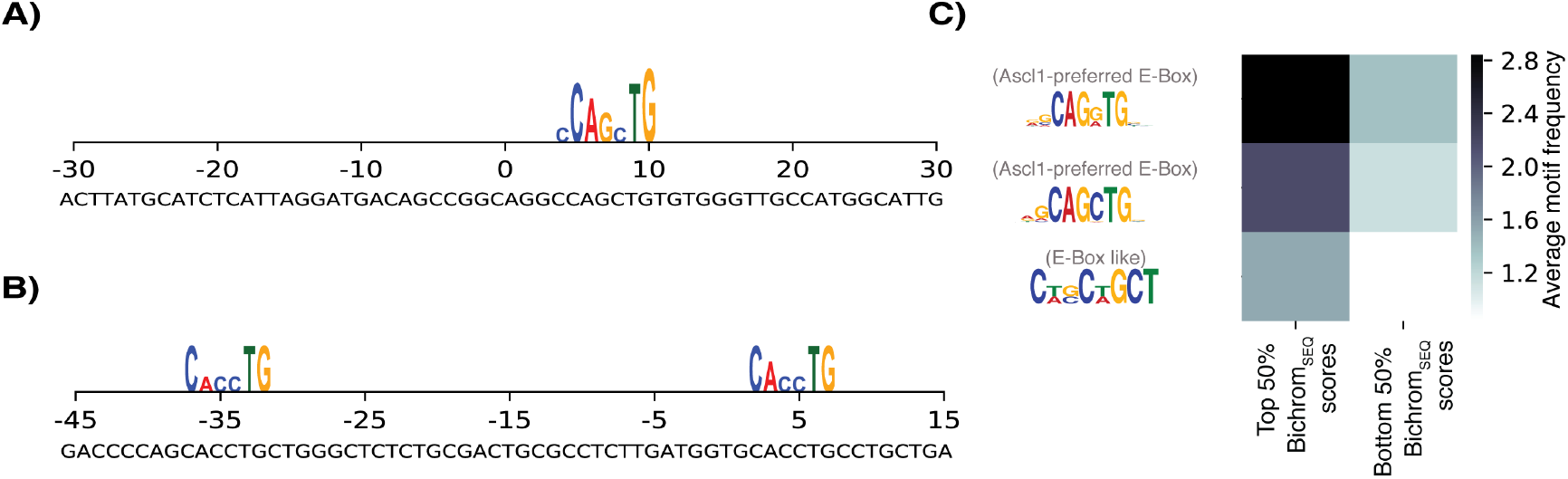
**A)** & **B)** Feature attribution with integrated gradients at two sample Ascl1 binding sites (chr10:4710120-4710170 and chr10:28136730-28136800). **C)** MEME-ChIP-determined E-box motif frequency at Ascl1 binding sites assigned the bottom 50% Bichrom_SEQ_ sub-network scores compared to sites assigned the top 50% Bichrom_SEQ_ sub-network scores.

**Supplementary Figure 7:**
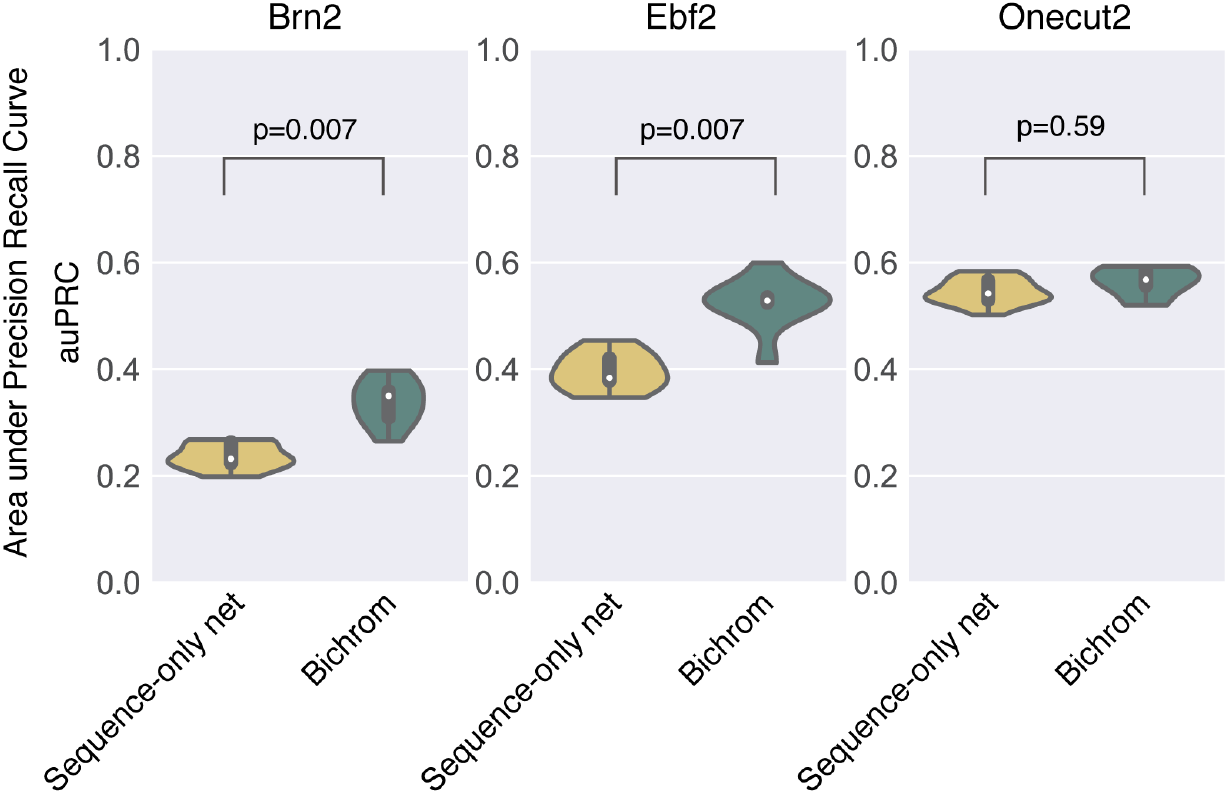
auPRC distributions for sequence-only and Bichrom networks trained on 9 independent training sets, each comprising of a distinct chromosome as a held-out test set for Brn2, Ebf2 and Onecut2.

**Supplementary Figure 8:**
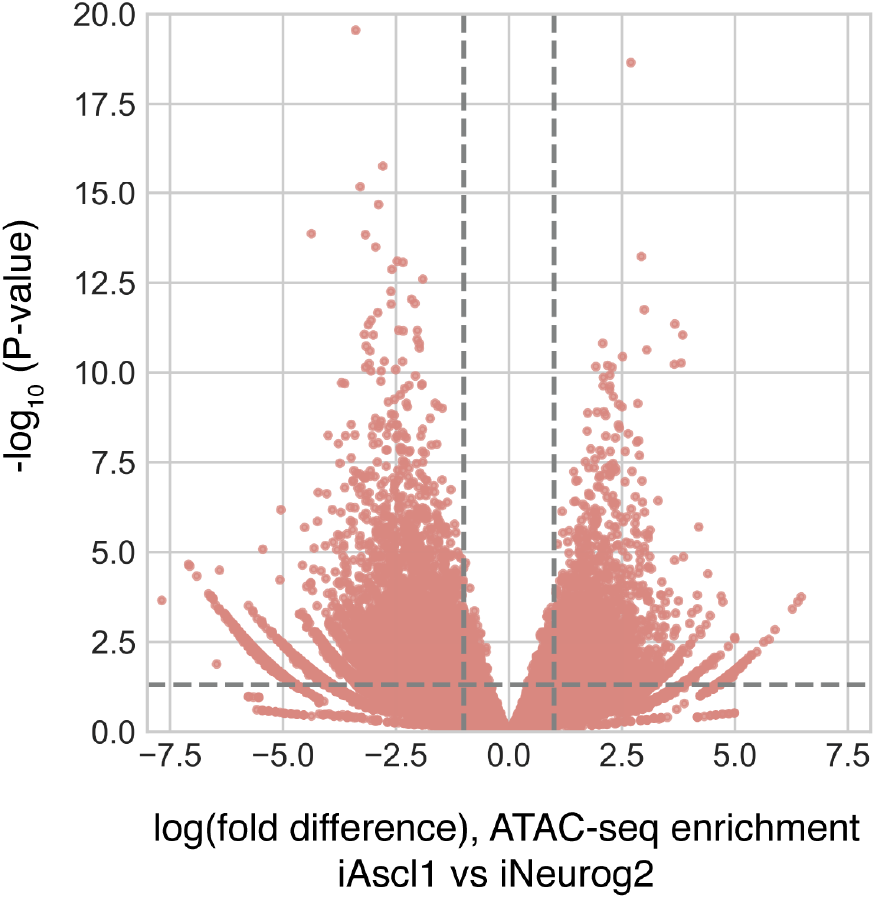
Differences in chromatin accessibility (ATAC-seq) levels in iAscl1+12hrs compared to iNeurog2+12hrs, as assessed using DESEQ2. The volcano plot maps the log fold-change and p-values at the union of accessible domains in iAscl1 and iNeurog2 neurons.

**Supplementary Figure 9:**
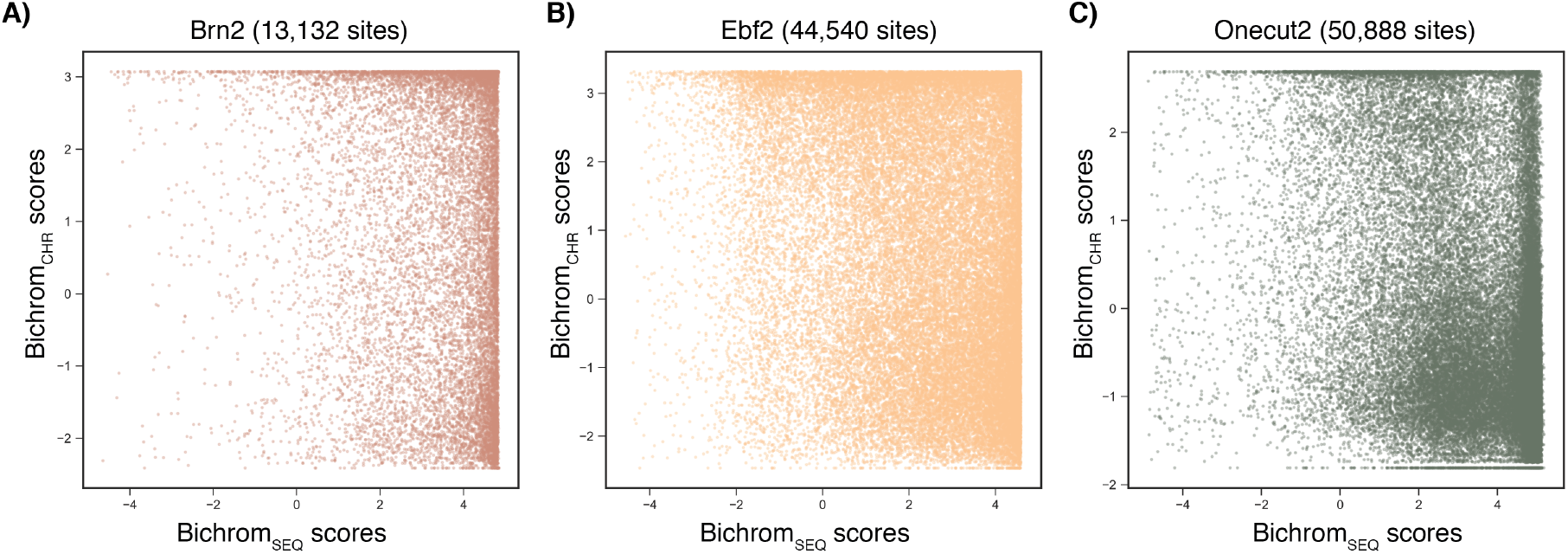
Bichrom-derived latent embeddings for **A)** Brn2, **B)** Ebf2 and **C)** Onecut2 induced in EB-derived iAscl1 cells.

**Supplementary Figure 10:**
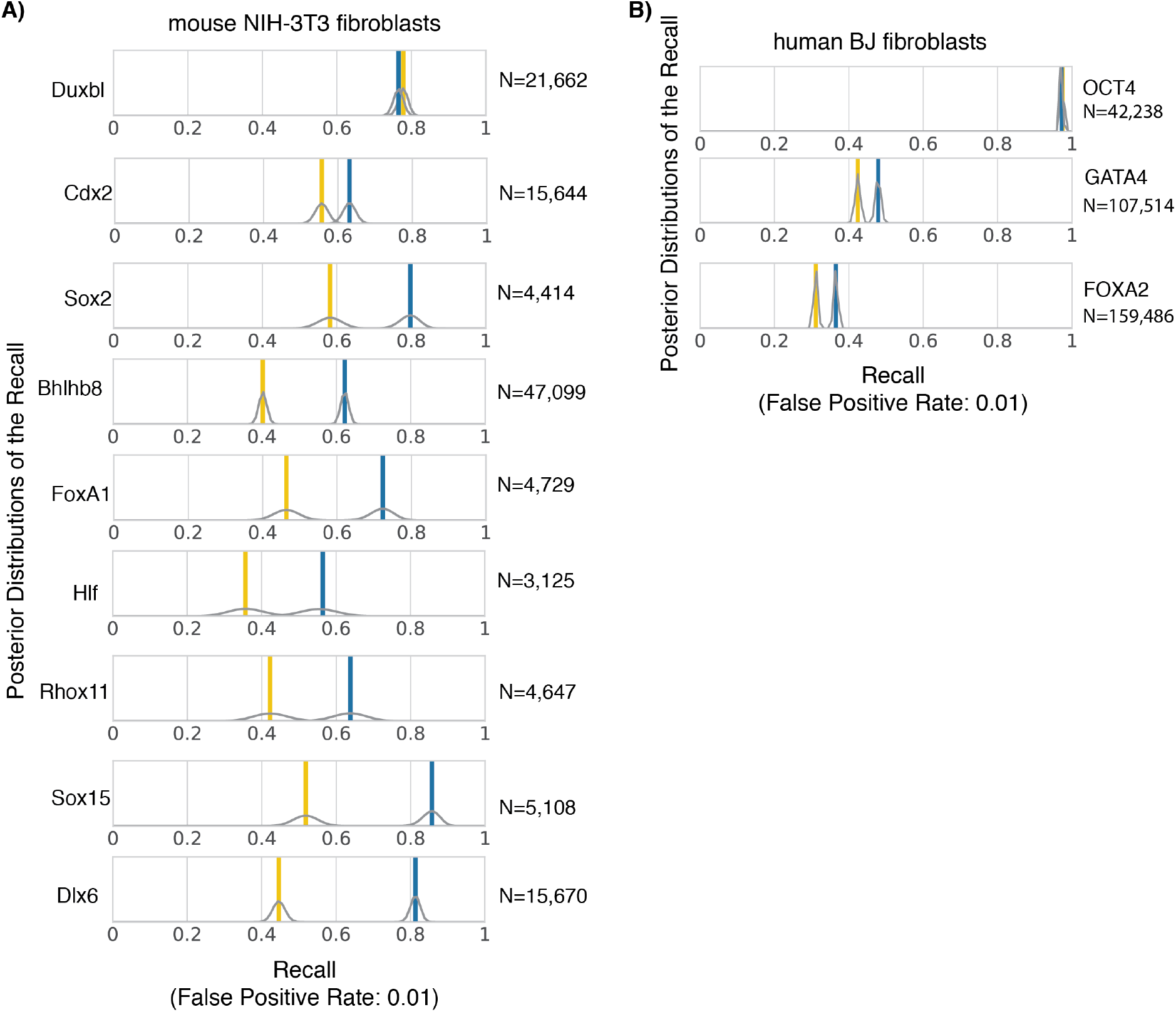
Bichrom recall at a fixed false positive rate (0.01) for **A)** the 9 mouse NIH-3T3 induced TFs and **B)** the 3 human fibroblast induced TFs. Chromosome 17 is used as a validation set, chromosome 11 is used as a held-out test set. The posterior distributions of the recall (estimated using a Beta uniform prior) are also plotted.

**Supplementary Figure 11:**
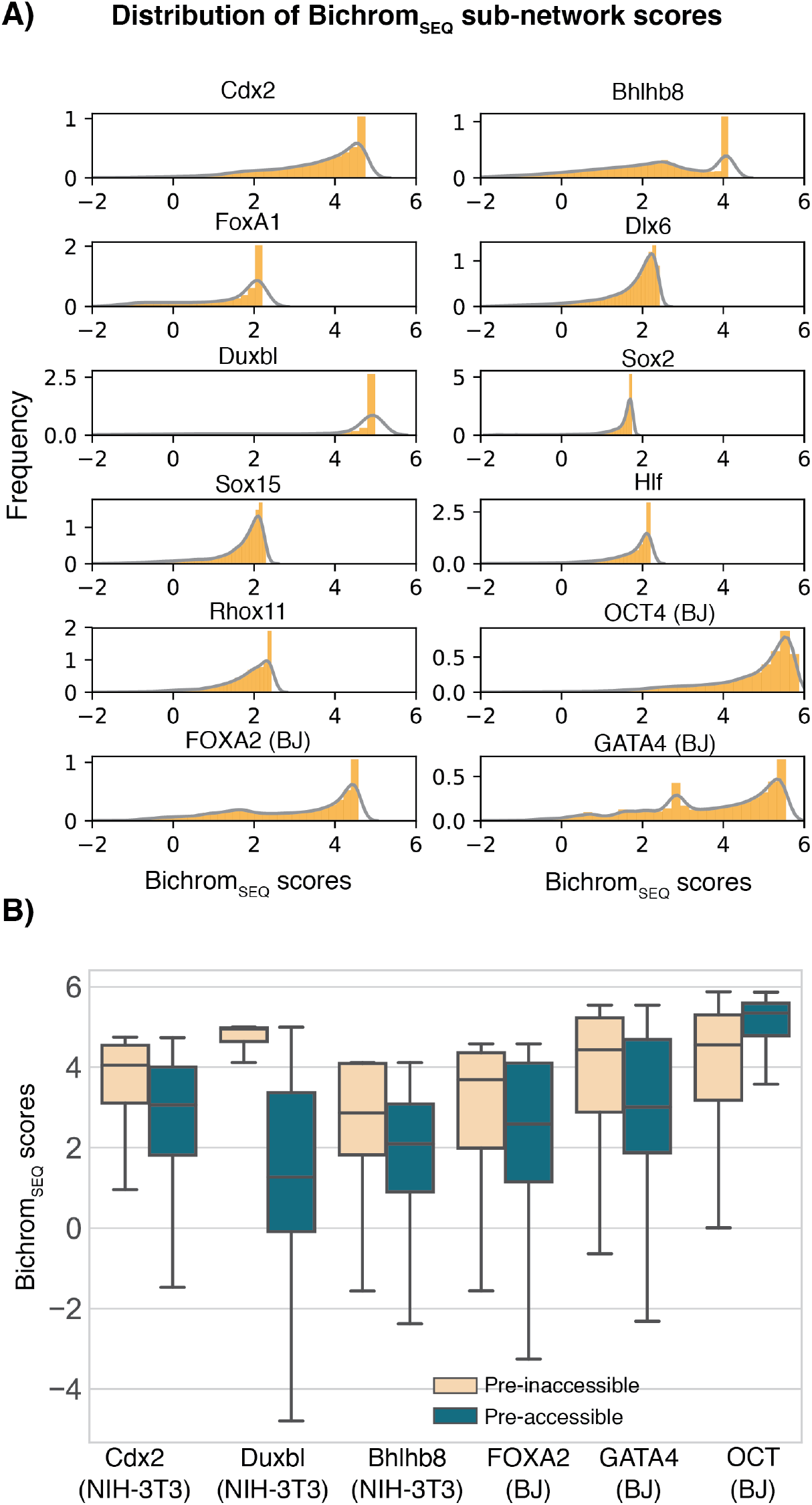
**A)** The distribution of Bichrom_SEQ_ sub-network scores for 9 TFs induced in mouse NIH-3T3 fibroblasts and 3 TFs induced in human BJ fibroblasts. **B)** The distributions of Bichrom_SEQ_ sub-network scores at pre-accessible versus pre-inaccessible binding sites are plotted for Cdx2, Duxbl, Bhlhb8, FOXA2, GATA4 and OCT4.

**Supplementary Figure 12:**
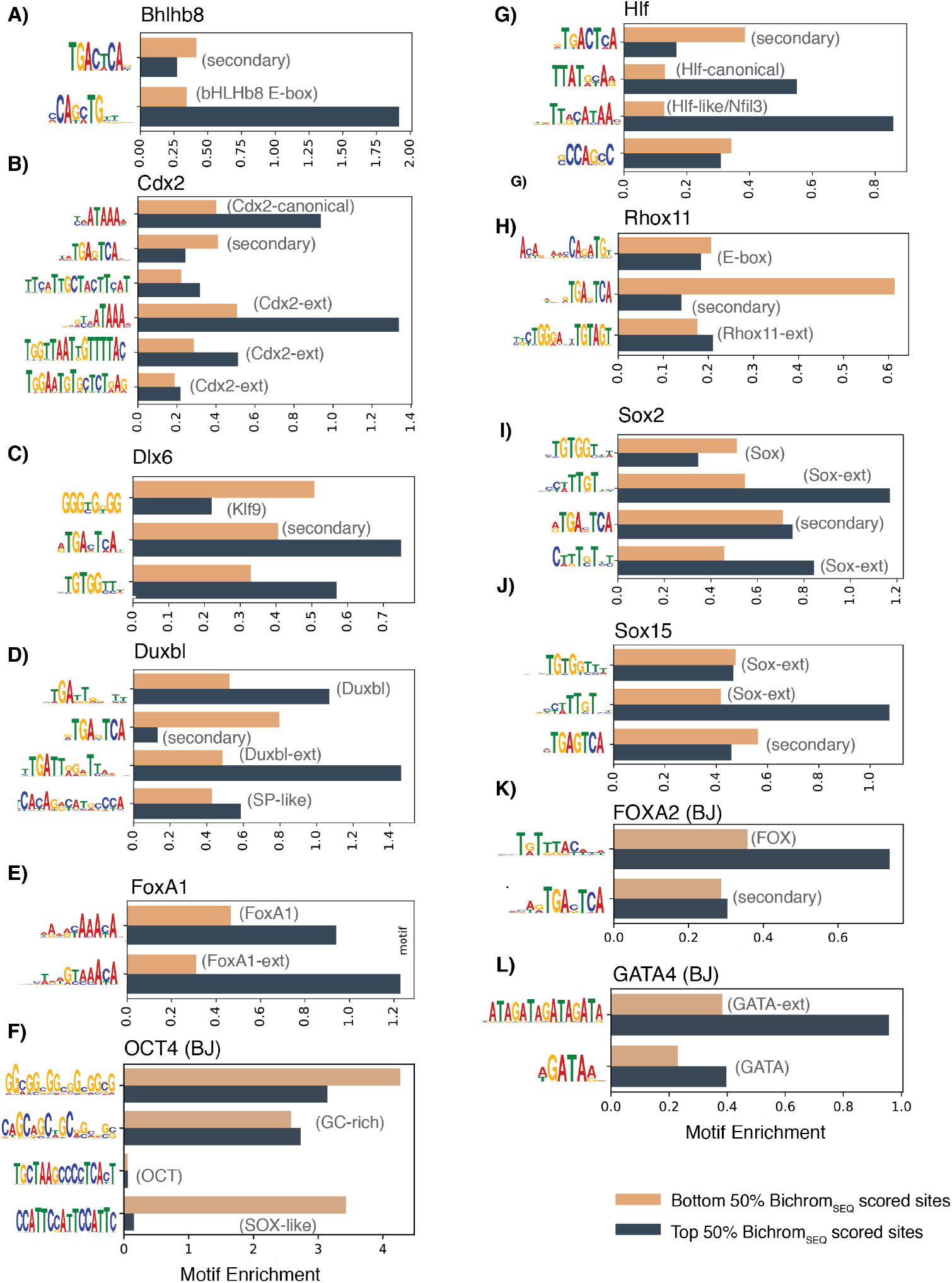
Analysis of MEME-ChIP-determined motifs for each induced TF. Individual plots show motif enrichment at sites assigned the bottom 50% Bichrom_SEQ_ sub-network scores compared to motif enrichment at sites assigned the top 50% Bichrom_SEQ_ sub-network scores for **A)** bHLHb8, **B)** Cdx2, **C)** Dlx6, **D)** Duxbl, **E)** FoxA1, **F)** OCT4 (BJ), **G)** Hlf, **H)** Rhox11, **I)** Sox2, **J)** Sox15, **K)** FOXA2 (BJ) and **L)** GATA4 (BJ).

**Supplemental Table 1:** Prior chromatin datasets for mouse embryoid bodies, generated in the same cell line as that were used to induce Ascl1.

| **Data** | **Replicates** | **Cell Type** | **Cell Line** |
| --- | --- | --- | --- |
| ATAC-seq<sup>1</sup> | 2 | EB | Ainv15 |
| H3K27ac<sup>1</sup> | 2 | EB | Ainv15 |
| H3K27me3<sup>1</sup> | 2 | EB | Ainv15 |
| H3K4me1<sup>1</sup> | 2 | EB | Ainv15 |
| H3K4me2<sup>1</sup> | 2 | EB | Ainv15 |
| H3K4me3<sup>1</sup> | 2 | EB | Ainv15 |

**Supplemental Table 2:** Prior chromatin datasets from mouse embryonic stem cells, sourced from the referenced publications.

| **Data** | **Replicates** | **Cell Type** | **Cell Line** |
| --- | --- | --- | --- |
| H3K9ac<sup>2</sup> | 2 | ES | E14 |
| H3K9me3<sup>2</sup> | 2 | ES | E14 |
| H3K36me3<sup>2</sup> | 2 | ES | E14 |
| H2A.Z<sup>3</sup> | 1 | ES | V6.5 |
| acH2A.Z<sup>3</sup> | 1 | ES | V6.5 |
| H4K20me3<sup>4</sup> | 2 | ES | V6.5 |

**Supplemental Table 3:**
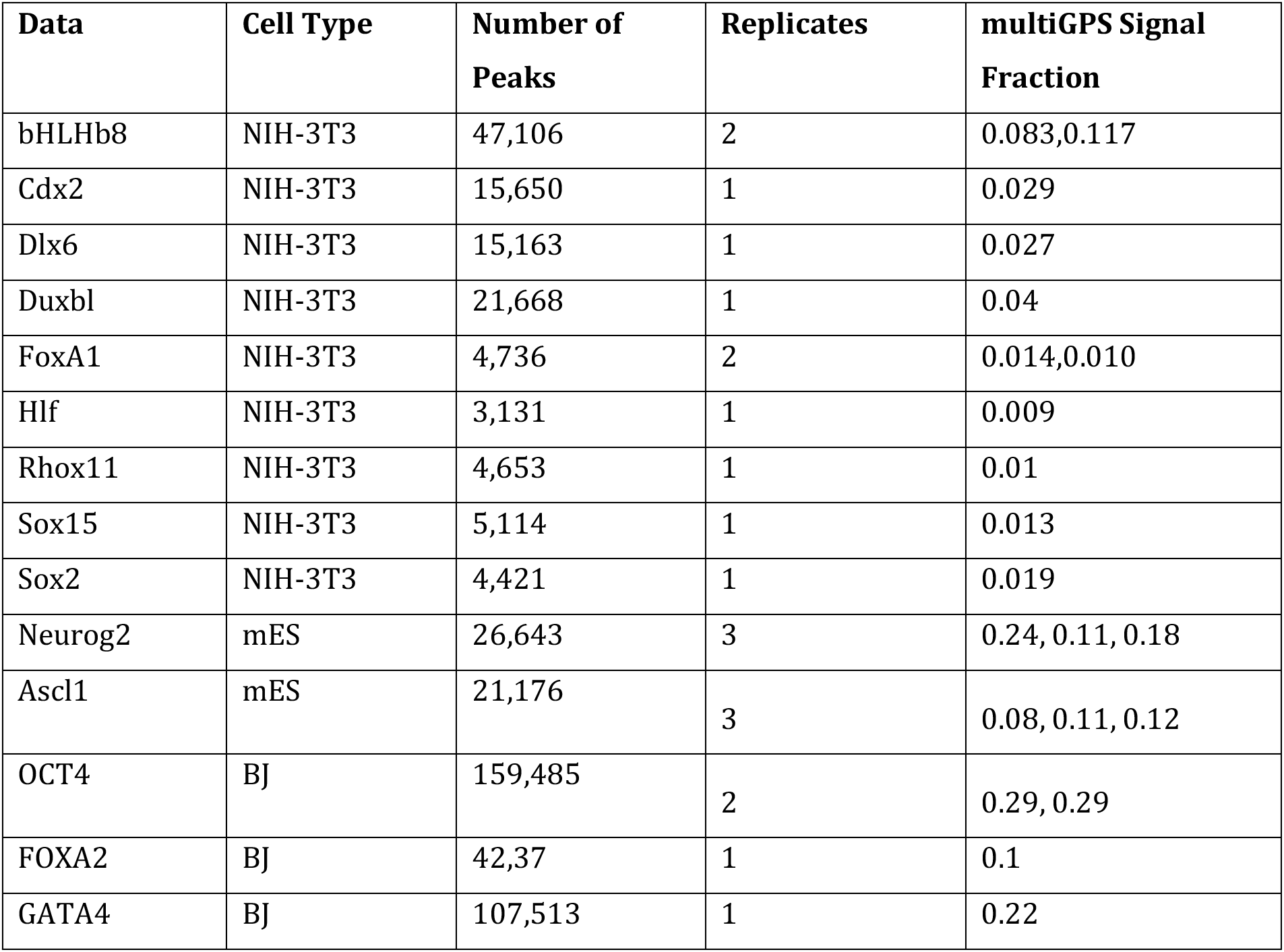
ChIP-seq data for TFs induced in NIH3T3 fibroblasts, downloaded from Raccaud *et al*^5^, mEB induced ChIP-seq data downloaded from Aydin *et al*^6^ and human BJ induced data, downloaded from Donaghey *et al*^7^. Processed with bowtie (1.0.1) and multiGPS (version 0.74).

**Supplemental Table 4:** ATAC-seq experiments used as prior chromatin data in mouse NIH3T3 fibroblasts^5^.

| Data | Replicates | Cell Type | Cell Line |
| --- | --- | --- | --- |
| ATAC-seq | 2 | Mouse Fibroblasts | NIH3T3 |

**Supplemental Table 5:** Experiments used as prior chromatin data in human BJ fibroblasts^7^.

| Data | Replicates | Cell Type | Cell Line |
| --- | --- | --- | --- |
| ATAC-seq | 2 | Human Fibroblasts | BJ |
| H2K27ac | 2 | Human Fibroblasts | BJ |
| H3K4me2 | 2 | Human Fibroblasts | BJ |
| H3K27me3 | 2 | Human Fibroblasts | BJ |

**Supplemental Table 6:** Bichrom network parameters tested within the hyper-parameter random grid search.

| Parameter | Options |
| --- | --- |
| Number of convolutional filters | 16, 32, 64, 128, 256 |
| Number of dense layers | 2-5 |
| Size of dense layers | 128, 512, 1024, 2048 |
| Activation functions | tanh, relu, sigmoid |
| Dropout Rate | 0.5, 0.25 |

**Supplemental Table 7:**
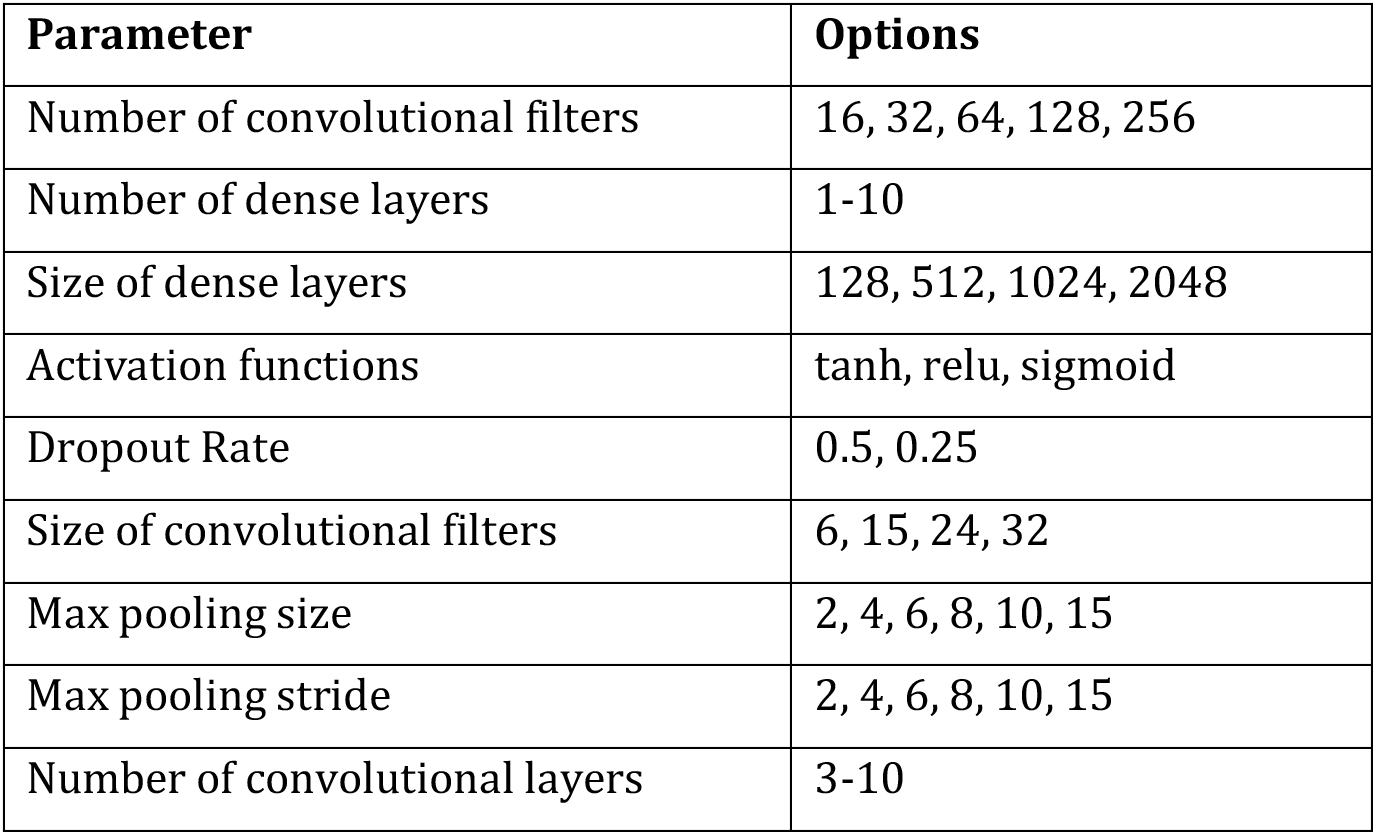
Network parameters used to construct alternative CNN-based networks (which are compared with the CNN-LSTM in Supp. Figure 1)

| Parameter | Options |
| --- | --- |
| Number of convolutional filters | 16, 32, 64, 128, 256 |
| Number of dense layers | 1-10 |
| Size of dense layers | 128, 512, 1024, 2048 |
| Activation functions | tanh, relu, sigmoid |
| Dropout Rate | 0.5, 0.25 |
| Size of convolutional filters | 6, 15, 24, 32 |
| Max pooling size | 2, 4, 6, 8, 10, 15 |
| Max pooling stride | 2, 4, 6, 8, 10, 15 |
| Number of convolutional layers | 3-10 |

